# Xist expression impacts chromatin structure and YY1 mediated transcription of X-linked genes in hematopoietic progenitor cells

**DOI:** 10.1101/2022.02.16.480654

**Authors:** Tianqi Yang, Jianhong Ou, Eda Yildirim

**Affiliations:** Department of Cell Biology, Duke University Medical Center; Department of Pharmacology and Cancer Biology, Duke University Medical Center; Duke Regeneration Center, Duke University; Duke Cancer Institute, Duke University Medical Center Durham, NC 27710 USA

**Keywords:** X chromosome inactivation, Xist RNA, epigenetic gene regulation, chromatin structure, transcription, hematopoiesis

## Abstract

X chromosome inactivation (XCI) is a dosage compensation phenomenon that occurs in females. Initiation of XCI depends on Xist RNA, which triggers silencing of one of the two X chromosomes, except for genes that escape XCI. This inactive state is propagated during cell divisions with continuous Xist expression. How Xist impacts XCI maintenance remains an open question. Here, we delete Xist in hematopoietic cells of mice and report increased fraction of cycling hematopoietic progenitors and differentiation defects during hematopoiesis. Xist loss results in upregulation of a subset of X-linked genes, several of which escape XCI and have functions in hematopoiesis and immunity. We find that the underlying mechanism of transcriptional upregulation involves reduced H3K27me3 occupancy and increased chromatin accessibility which accommodate binding of YY1 transcription factor (TF). We propose that XCI maintenance defects in Xist deficient mice rely on the landscape of open chromatin sites, TF availability, and cell cycle stress in specific cellular context during hematopoiesis.

## INTRODUCTION

X chromosome inactivation (XCI) is a dosage compensation phenomenon that occurs in mammalian females to resolve the inherent X-linked gene dosage imbalance between sexes through transcriptional silencing one of the two X chromosomes ^1^. The random form of XCI occurs in the epiblast on embryonic days 4.5-5.5 (E4.5-5.5). This phase is referred as the ‘XCI initiation’ phase during which either X chromosome has the equal probability to be chosen as the future inactive X chromosome (Xi). With the establishment of the transcriptionally silent state, the cells enter into a “XCI maintenance” phase in which the same X chromosome is propagated as the Xi in subsequent cell divisions. Proper maintenance of the Xi silent state is essential for female health because XCI maintenance defects have long been implicated in various human cancers and development diseases ^1–6^. Therefore, determining the molecular basis of XCI maintenance is important to provide insights towards development of therapies for X-linked diseases.

Initiation of XCI depends on Xist RNA which is a 17-kb transcript that is exclusively expressed from the Xi ^7, 8^. Xist RNA binds across the Xi in *cis*, and aids recruitment of chromatin regulatory factors including the Polycomb Repressive complexes (PRC1 and PRC2) ^9–11^ and YY1 ^12^ that influence the establishment of the silent chromatin state on the Xi. These chromatin changes lead to heterochromatinization across the Xi with enrichment of repressive histone marks including Histone 3 Lysine 27 tri-methylation (H3K27me3) ^13^. In addition to the silent chromatin features, the Xi adopts a unique higher chromatin organization with low chromatin accessibility ^14, 15^. In contrast to the majority of the X-linked genes, genes that escape XCI continue to be biallelically expressed ^16, 17^. These genes exhibit open chromatin accessibility and associate with enriched active histone modification, H3K4me3, at transcription start sites (TSS) and low H3K27me3 across gene bodies ^18, 19^.

Expression of Xist during the initiation of XCI is essential for embryonic development ^20^. Xist is continuously expressed in the *soma* during the lifetime of the female. However, Xist has been thought to be dispensable for the maintenance of silencing based on studies that utilized embryonic stem cells and fibroblasts ^21–23^, and until recently only a few *in vivo* studies have been reported suggesting a role for Xist in XCI maintenance ^20, 24, 25^. We contributed to these earlier studies by showing that Xist deletion in hematopoietic stem cells (HSCs) leads to progressive upregulation of X-linked genes suggesting a potential role for Xist in XCI maintenance ^25^. We found that Xist loss results in altered hematopoiesis with expansion of hematopoietic stem and progenitor cells (HSCPs), and differentiation and maturation abnormalities in all three major cell lineages including myeloid, erythroid and lymphoid cells ^25^. Notably, Xist mutant mice exhibit female-specific lethality due to a range of aggressive hematologic malignancies in the form of mixed myelodysplastic syndrome/myeloproliferative neoplasm (MDS/MPN) and chronic myelomonocytic leukemia (CMML). Similar to the hematopoietic system, targeting of Xist in other cell systems including skin, gut, brain, and the epiblast have revealed that Xist loss leads to partial upregulation of X-linked genes, yet unlike the hematopoietic system many tissues exhibit high-tolerance to Xist loss ^26, 27^. These findings suggest variable functions of Xist in different tissues, and argue that the hematopoietic system provides an excellent system to study undiscovered functions of Xist during XCI maintenance.

Here, we conditionally deleted Xist in the HSCs of mice ^25^ to determine the functional relevance of Xist during hematopoiesis. At the cellular level, we found that Xist loss led to significant changes in the ratio of hematopoietic progenitor cells and increase in the fraction of cycling pluripotent progenitor cells. At the molecular level, we identified ’lineage-specific’ and ’common’ differentially expressed genes (lsDEGs and cDEGs) in different Xist deficient HSPC populations. We determined that the upregulated X-linked cDEGs were enriched in XCI escape genes ^16^. Several of these escape genes have been functionally associated with hematopoiesis and immune response suggesting that their upregulation might be causally linked to the development of systematic hematopoietic defects in Xist deficient female mice. By analyzing changes in the distribution of histone modifications and chromatin accessibility, we further revealed that the underlying mechanism of transcriptional upregulation of a subset of X-linked cDEGs involve reduced H3K27me3 occupancy, increased chromatin accessibility, and enrichment in YY1 binding. Collectively, our findings provide insights on how proper Xist expression during XCI maintenance influences X-linked gene regulation at the level of chromatin accessibility, histone modifications and binding of TFs, and offer a potential explanation to why hematopoietic cells exhibit sensitivity to Xist loss on the basis of their cell cycle regulation. Findings of this study are critical towards identification of novel targets for therapy for X-linked diseases.

## RESULTS

### Lineage specific differentiation defects during hematopoiesis in Xist deficient mice

Deletion of Xist in HSCs leads to progressive hematopoietic defects resulting in hematologic malignancies and lethality in female mice ^25^. In contrast, deletion of Xist in more differentiated cells such as pro-B lymphocytes does not result in any obvious illness or lethality ^28^. To investigate the functional role of Xist in HSPCs, we utilized mice at early disease stages (up to 5 month-old) and quantified the number and percentage of different HSPC populations by flow cytometry ^29^ (Figure 1A and S1A). We focused on three HSPC populations in the bone marrow, including LSK+ (Lin-Scal1+c-Kit+) cells, LSK-(Lin-Scal1-c-Kit+) cells, and Lin-(lineage-negative) cells (Figure 1B). Consistent with our previous findings ^25^, we observed a significant increase in the percentage (∼2.5 fold; *p*<0.01) and number (∼2 fold; *p*<0.001) of bone marrow LSK+ cells of female XistΔ/Δ mice in comparison to their wild-type (WT) littermates (Figure 1A, 1C-D). LSK+ cells are composed of four subpopulations that can be characterized using SLAM cell-surface markers ^30^. These subgroups include HSCs (Lin^-^c-Kit^+^Sca-1^+^CD150^+^CD48^-^), multipotent progenitor cells (MPPs) (Lin^-^c-Kit^+^Sca-1^+^CD150^-^CD48^-^) and two hematopoietic progenitor populations with restricted pluripotency, HPC1 (Lin^-^c-Kit^+^Sca-1^+^CD150^-^CD48^+^) and HPC2 (Lin^-^c-Kit^+^Sca-1^+^CD150^+^ CD48^+^) (Figure 1A-B, and S1A). We found significant expansion in the number of HPC1 (∼2.5-fold; *p*<0.001) and HPC2 (∼3-fold; *p*<0.001) cells in XistΔ/Δ female mice (Figure 1A and 1D).

**Figure 1:**
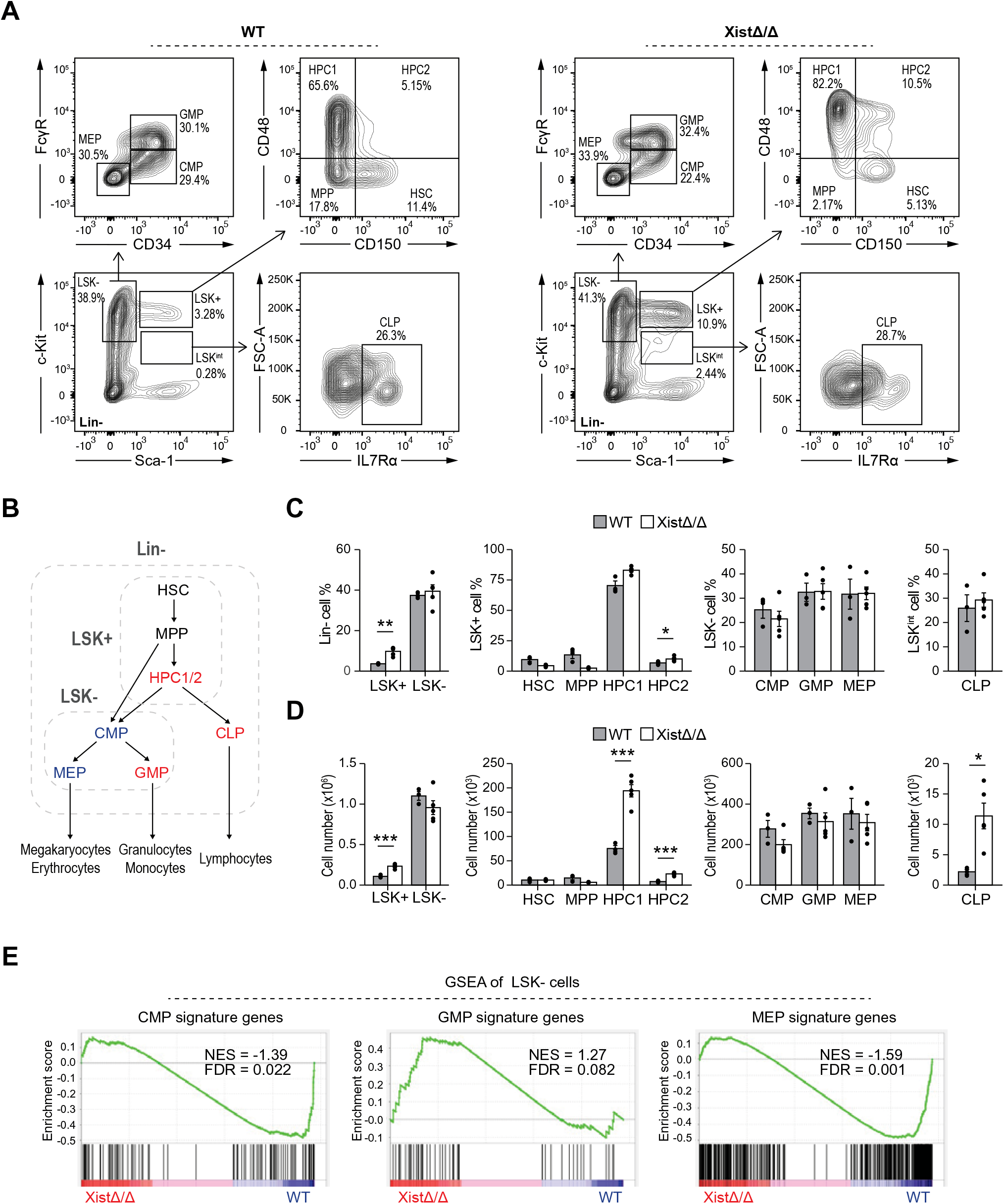
Lineage-specific differentiation defects of hematopoietic progenitors in Xist deficient mice. **(A)** Representative FACS analysis of Lin-bone marrow cells from a 3-month-old XistΔ/Δ and WT female mice. **(B)** Schematic shows differentiation hierarchy of HSPCs. Cell populations that exhibit increase or decrease in cell number or signature gene expression in Xist deficient female mice are highlighted in red and blue, respectively. **(C and D)** FACS analysis of LSK+, LSK- and their subpopulations show increased percentage of LSK+ cells and HPC2 cells (C), and increased number of LSK+, HPC1, HPC2 cells, as well as increased number of LSK- CLP cells (D) in 3-month-old XistΔ/Δ (n=5) and WT (n=3) female mice. Means ± SEM are shown; Student’s *t*-test, ***p <0.001, **p<0.01, *p<0.05. **(E)** Gene set enrichment analysis (GSEA) show significant decrease in the expression of CMP- and MEP signature genes and slight increase in GMP signature genes in 3-month-old XistΔ/Δ in comparison to WT female mice. Normalized enrichment scores (NES) and false discovery rates (FDR) are indicated in corresponding plots. FDR < 0.05 is defined as significant. CMP, common myeloid progenitor; MEP, megakaryocyte-erythroid progenitor cell; GMP, granulocyte-monocyte progenitors.

To investigate whether expansion of HPC1/2 cells affected downstream lineage-specific progenitors, we stained Lin- bone marrow cells using cell surface markers IL7Rα, CD34 and FcγR ^30^ (Figure 1A and S1A). Our FACS analyses revealed that the number of common lymphoid progenitors (CLP; Lin^-^ c-Kit^+^ Sca-1^int^IL7R*α*^+^) were significantly increased (∼5 fold; *p*<0.05) upon Xist deletion (Figure 1D). In contrast, the percentage or the number of common myeloid progenitors (CMP; Lin^-^c-Kit^+^Sca-1^-^CD34^+^Fc*γ*R^int^) or the two downstream myeloid progenitor populations, including granulocyte-monocyte progenitors (GMP; Lin^-^c-Kit^+^Sca-1^-^CD34^+^Fc*γ*R^+^) and megakaryocyte–erythroid progenitors (MEP; Lin^-^c-Kit^+^Sca-1^-^CD34^-^Fc*γ*R^-^) (Figure 1A, 1C-D) were comparable between Xist deficient and WT mice. To determine whether Xist deficiency influences transcription of LSK- cell subpopulations, we utilized RNA-Seq. We generated differential expression profile of LSK- cells in XistΔ/Δ and WT female mice and performed Gene Set Enrichment Analyses (GSEA) ^31^ with signature gene sets of LSK- subpopulations including CMP, GMP and MEP (See Methods for details of the analysis) (Figure S1B). Despite the lack of changes in the number of GMP, CMP and MEP (Figure 1C and 1D), we detected a slight transcriptional upregulation of GMP signature genes and significant reduction in the transcription of CMP (FDR=0.022) and MEP (FDR=0.001) signature genes in XistΔ/Δ LSK- cells (Figure 1E). We concluded that Xist deletion facilitates lymphoid progenitor differentiation and hinders myeloid progenitor differentiation, specifically towards the megakaryocyte and erythrocyte lineage.

### Increased fraction of cycling hematopoietic progenitor cells in Xist deficient mice

Differentiation and proliferation of HSPCs are finely controlled by cell cycle checkpoints ^32^. Given the significant changes in the number of LSK+ cells, we investigated how Xist loss impacts cell cycle in LSK+ and LSK- cells by FACS (see schematic of the experiment design in Figure 2A (left)). Cells were stained for the cell proliferation marker, Ki67, to discriminate quiescent cells at G0 phase (Ki67^-^) and cycling cells that are at G1/S/G2/M phases (Ki67^+^) (Figure 2A). In WT mice, quiescent cells constituted ∼90% of the LSK+ cells, and ∼45% of the LSK- cells (Figure 2A-B). In XistΔ/Δ female mice, we detected a significant increase in the percentage of cycling MPP (∼3-fold; *p*<0.05), HPC1 (∼3-fold; *p*<0.01) and HPC2 (∼2-fold; *p*<0.05) cells (Figure 2A, 2B and S1C). In contrast, Xist mutant HSCs and LSK- cells showed comparable cell cycle patterns with WT cells (Figure 2A, 2B, and S1C).

**Figure 2:**
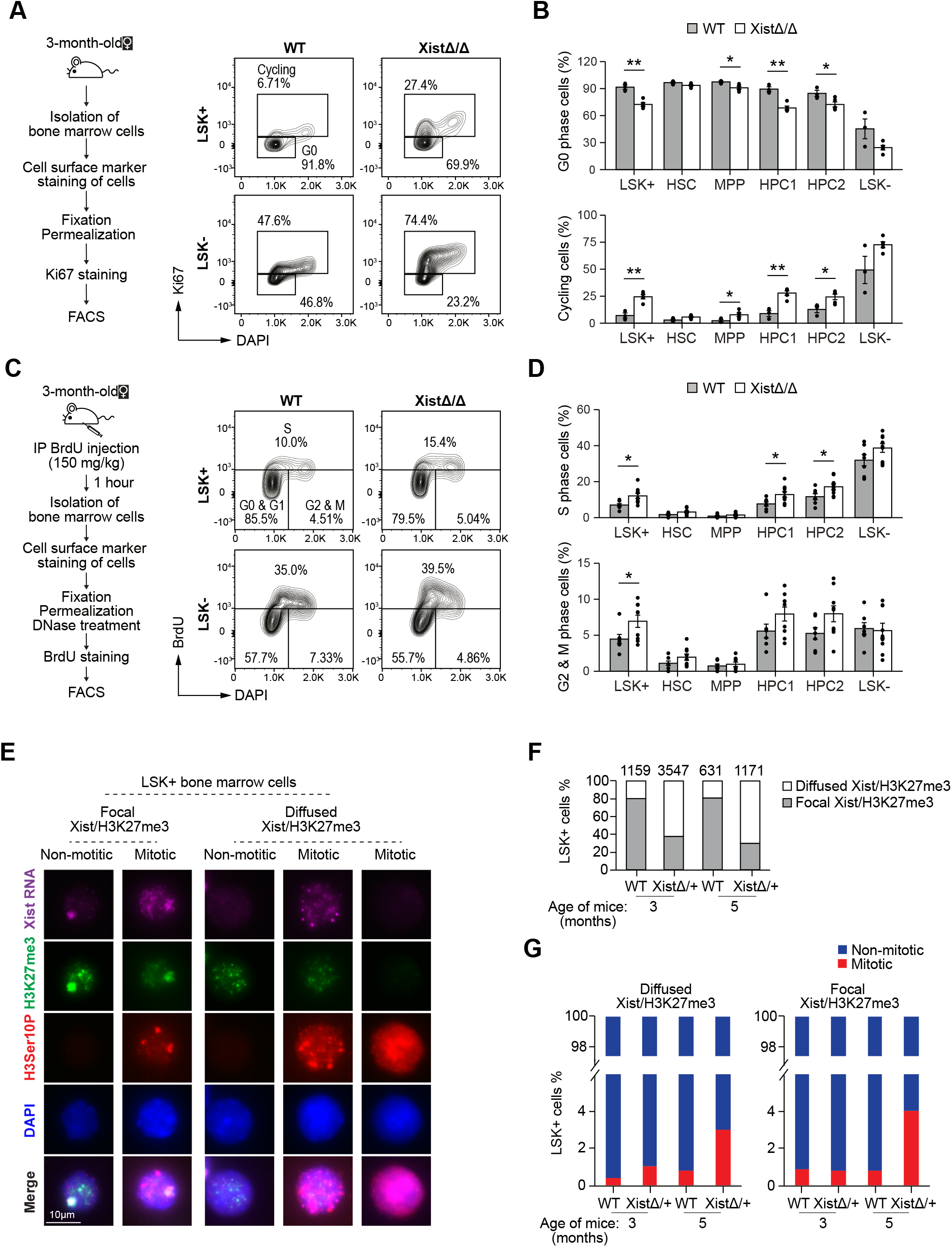
Increased faction of cycling hematopoietic progenitor cells in Xist deficient mice. **(A)** Schematic shows experimental design of FACS based Ki67 cell cycle assay (left); Representative FACS analysis of bone marrow cells from a 3-month-old female XistΔ/Δ and WT female littermate mouse (right). **(B)** FACS analyses show significant increase in the percentage of cycling and decrease in the percentage of quiescent LSK+, MPP, HPC1, and HPC2 cells in 3-month-old XistΔ/Δ (n=5) in comparison to WT (n=3) female mice. Means ± SEM are shown; Student’s *t*-test, **p<0.01, *p<0.05. **(C)** Schematic shows experimental design of FACS based BrdU cell cycle assay (left); Representative FACS analysis of bone marrow cells from a 3-month-old female XistΔ/Δ and its WT female littermate mouse (right). **(D)** FACS analyses show significant increase in the percentage of LSK+, HPC1, HPC2 cells at S- phase, and LSK+ cells at G2/M phase in 3-month-old XistΔ/Δ (n=9) in comparison to WT (n=7) female mice. Means ± SEM are shown; Student’s *t-*test, **p<0.01, *p<0.05. **(E)** Representative images of sequential Xist RNA FISH and IF staining of H3K27me3 and H3Ser10P using WT LSK+ bone marrow cells show focal and diffused Xist and H3K27me3 signals. Scale bar, 10 μm. **(F)** Analysis of cells with focal or diffused Xist/H3K27me3 IF staining shows decrease in the percentage of LSK+ cells that show focal Xist/H3K27me3 signal in 3-month-old Xist+/Δ female mouse in comparison to WT female mouse. The affect becomes more pronounced with age when 5-month-old mice are examined. Number of analyzed LSK+ cells are as shown in the bar graph. **(G)** Analysis of mitotic and non-mitotic cells in LSK+ cells with diffused Xist/H3K27me3 (left) or focal Xist/H3K27me3 (right) show progressive increase in the percentage of LSK+ mitotic cells in 3- and 5-month-old Xist+/Δ female mouse in comparison to WT female mouse. In 5-month-old Xist+/Δ mouse, the percentage of mitotic LSK+ cells was increased independent of the focal or diffused Xist/H3K27me3 signal. Number of analyzed LSK+ cells are as shown in the bar graph in Figure 2F.

We next performed in vivo cell cycle analysis by labeling bone marrow cells with BrdU ^33^ (see schematic of the experiment design in Figure 2C (left)). Consistent with the Ki67-based cell cycle analyses, we detected a significant increase in the percentage of Xist deficient LSK+ (∼1.7 fold; *p*<0.05), HPC1 (∼1.7 fold; *p*<0.05) and HPC2 (∼1.5 fold; *p*<0.05) cells at the S-phase (Figure 2C, 2D, and S1C). These data supported increased cell cycle activity specifically in hematopoietic progenitors with restricted pluripotency in Xist deficient female mice.

Cytological analyses of the spleen and bone sections, and bone marrow smears revealed increased mitotic figures in Xist deficient mice ^25^. In support of this data, we found that the fraction of LSK+ cells at G2/M phase was significantly higher (∼1.6 fold; *p*<0.05) in Xist deficient female mice compared to their WT counterparts (Figure 2D). These data prompted us to investigate mitotic cells at single cell resolution. We performed sequential Xist RNA FISH and immunofluorescence (IF) against H3K27me3 and mitotic marker, Histone 3 Serine 10 phosphorylation (H3Ser10P), in LSK+ cells (Figure 2E). In line with earlier reports ^24^, ∼80% of WT LSK+ cells showed a single focal Xist RNA cloud on the Xi that overlapped with H3K27me3 staining (Figure 2F). As expected, the percentage of cells with focal Xist/H3K27me3 staining was reduced almost by half in XistΔ/+ female mice at early disease stage (3-month-old), and such an effect became slightly more pronounced at later disease stage (5-month-old) (Figure 2F). We next quantified the percentage of mitotic LSK+ cells that showed diffused or focal Xist/H3K27me3 staining between WT and Xist deficient mice. In 3-month-old mice, we detected a slight increase in the percentage of mitotic LSK+ cells with diffused Xist/H3K27me3 staining (Figure 2G). In contrast, we observed a drastic increase in the percentage of Xist deficient mitotic LSK+ cells with focal and diffused Xist/H3K27me3 staining in 5-month-old mice (Figure 2G). Based on these findings, we concluded that there is a progressive increase in the cell cycle activity of Xist deficient LSK+ cells during progression of hematopoietic malignancies in Xist mutant mice, and that such change relies on both primary effects of Xist loss and the secondary effects of systematic hematopoietic stress that accumulate over time.

### Variable regulation of X-linked gene expression in Xist deficient hematopoietic cells

Previous microarray based transcriptional profiling of multiple hematopoietic cell populations in Xist deficient female mice have shown that transcription of several X-linked genes was progressively upregulated during disease progression^25^. Here, we utilized RNA-Seq and generated a comprehensive transcription profile for Lin-, LSK+ and LSK- cells in WT and Xist deficient mice (n=2/genotype). Cumulative distribution fraction (CDF) analysis of expression fold changes of all transcriptionally active genes (FPKM ≥ 1) revealed a slight, but significant right shift of X-linked curve compared to the autosomal curve in Xist deficient LSK+ cells (p=0.03137) (Figure 3A). Right shift detected in X-linked curve became more pronounced in Lin- Xist deficient cells (p=3.473e-05) (Figure 3A). This finding demonstrated that Xist deletion results in upregulation of X-linked gene expression and thus may lead to dosage compensation defects during differentiation of Xist-deficient cells. In contrast, the X-linked CDF curve was comparable to the autosomal curve in Xist deficient LSK- cells (Figure 3A), supporting our findings that myeloid and lymphoid progenitors exhibit variable tolerance to Xist loss (Figure 1).

**Figure 3:**
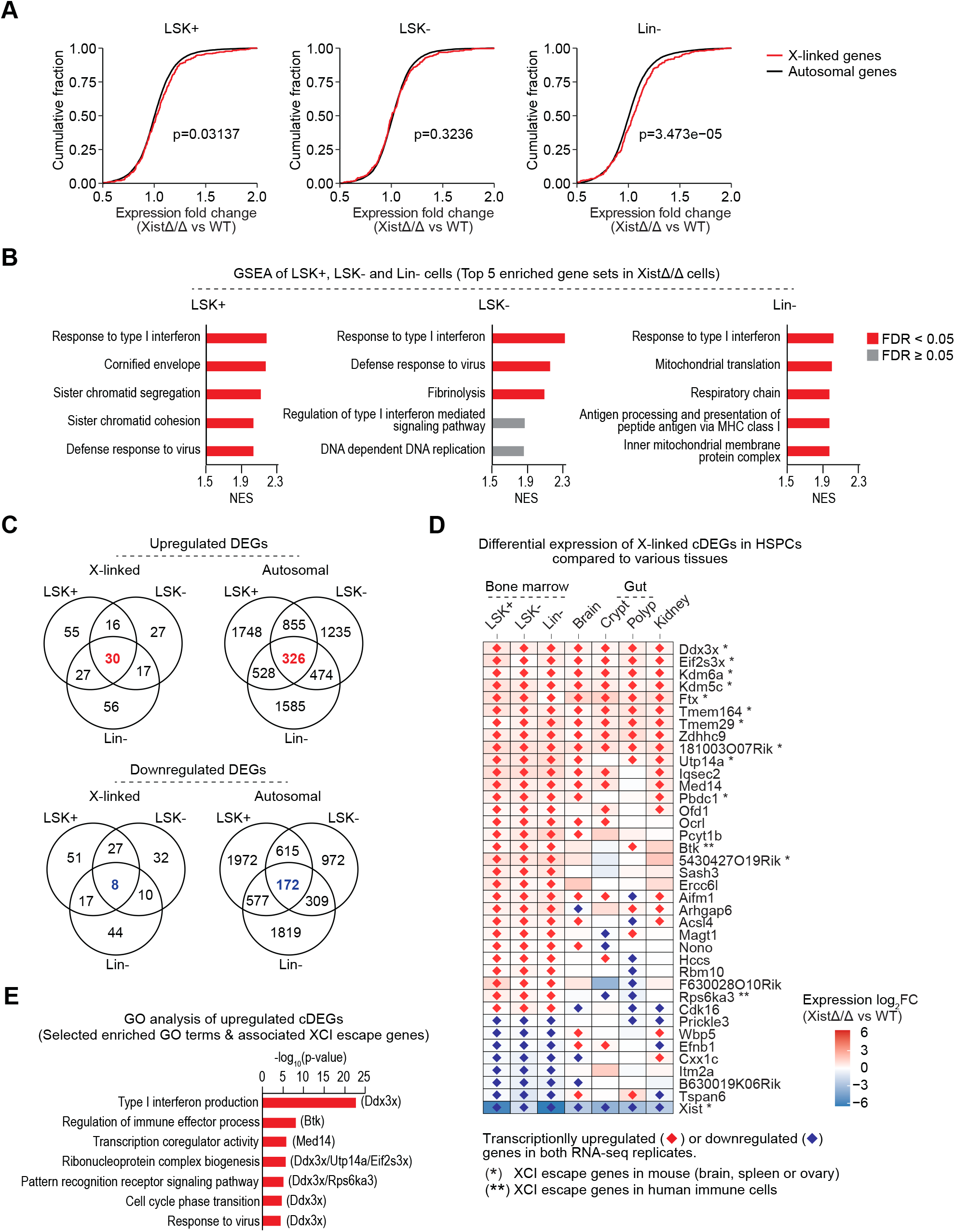
Transcriptional regulation of X-linked genes in Xist deficient hematopoietic cells. **(A)** Cumulative distribution fraction (CDF) plot of expressional fold change of X-linked and autosomal genes between XistΔ/Δ and WT mice show significant increase in expression of X-linked genes in XistΔ/Δ LSK+ and Lin- cells. Genes with FPKM>1 were used for plotting. Difference between X-linked and autosomal CDF curves was tested by Kolmogorov-Smirnov statistic and p-values are as indicated. **(B)** GSEA of LSK+, LSK- and Lin- cells show upregulated expression of genes related to Type I interferon-mediated immune responses in all three Xist-deficient HSPC populations, and genes related to cell division in XistΔ/Δ LSK+ cells. Top 5 gene sets enriched in XistΔ/Δ cells (ranked by NES) were plotted. See Table S1 for complete GSEA results. **(C)** Venn diagrams show overlap between differentially expressed X-linked or autosomal genes (DEGs) in LSK+, LSK- and Lin- cells. Upregulated or downregulated DEGs that are shared in all three cell types are highlighted in red and blue, respectively. See Table S1 for a full list of DEGs. **(D)** Heatmap shows expression of Log2FC of X-linked cDEG in bone marrow cells (LSK+, LSK- and Lin- cells) in comparison to various tissues including brain ^27^, gut (crypt and polyp) and kidney ^28^. Genes that were characterized as XCI escape genes in mouse ^34^ or human immune cells ^55, 56, 86^ are marked with an asterisks. **(E)** GO analysis show that biological function of transcriptionally upregulated cDEGs associate with Type I interferon production, immune response, cell cycle regulation and transcriptional regulation. Names XCI escape genes that associate with each GO term were indicated in the graph. For a complete list of GO analysis see Table S1.

We next analyzed differential expression profiles of LSK+, LSK- and Lin- cells by GSEA. Our data revealed that expression of genes that associate with Type-I interferon-mediated immune responses to virus infection were enriched in all three XistΔ/Δ HSPC populations (Figure 3B and Table S1). On the other hand, expression of genes related to chromatin segregation, DNA replication and cell division were enriched only in LSK+ cells compared to LSK- cells in Xist deficient mice (Figure 3B and Table S1). This data is in accordance with the results of our cell cycle analysis showing increased cell cycle activity in Xist deficient LSK+ cells (Figure 2). These results suggest that genes with functions in hematopoiesis, immune response and cell cycle might contribute to development of hematologic malignancies in Xist-deficient female mice.

To determine candidate genes that might be causally linked to functional defects in Xist-deficient cells, we focused on differentially expressed genes (DEGs) that show consistent transcriptional upregulation or downregulation in both biological replicates of a given cell lineage (Table S1). We found that majority of DEGs were expressed in one or two HSPC populations. We referred to these genes as “lineage-specific" differentially expressed genes or lsDEGs. In addition, hundreds of DEGs were expressed in all three HSPCs and we referred to these genes as “common" differentially expressed genes or cDEGs (Figure 3C and Table S1). Compared to the autosomal cDEGs (326 upregulated; 172 downregulated), X-linked cDEGs (30 upregulated; 8 downregulated) presented a strong bias for the transcriptionally upregulated genes (Figure 3C). Notably, almost one third of X-linked cDEGs (11/30) (Ddx3x, Eif2s3x, Kdm6a/Utx, Kdm5c/Jarid1c, Ftx, Tmem164, Tmem29, 1810030O07Rik, Utp14a, Pbdc1 and 5430427O19Rik) have been characterized as XCI escape genes in mouse brain, spleen, or ovary ^34^ (Figure 3D and Table S1). Importantly, most of these reported XCI escape genes also showed upregulation in other tissues with relatively high tolerance to Xist deletion (Figure 3D) ^27, 28^, suggesting that their upregulation was a direct effect of Xist loss rather than secondary responses to cellular stress. Collectively, this data indicated that Xist influences transcription of X-linked genes through lineage-dependent and -independent manner, the latter regulation preferentially targeting XCI escape genes.

Gene ontology (GO) analysis of upregulated cDEGs revealed that many of the upregulated X-linked cDEGs associated with biological processes including Type I interferon production, immune responses, cell cycle regulation, transcription regulation, and ribonucleoprotein complex biogenesis (Figure 3E and Table S1). Importantly, each of these biological processes associates with one or more reported XCI escape genes as highlighted in Figure 3E (Table S1). Overall, our transcriptional analysis suggested that a subset of X-linked genes, among which many have been reported to escape XCI, are highly susceptible to transcriptional upregulation upon Xist loss. Given the biological functions of these genes, we concluded that Xist expression is required for maintaining transcription of X-linked genes that associate with cell homeostasis and proper function of hematopoietic cells in female organism.

### Altered distribution of H3K27me3 across the X chromosome does not exclusively correlate with transcriptional upregulation of X-linked genes in Xist deficient cells

Recruitment of the PRC2 onto the Xi is one of the early XCI initiation events that triggers heterochromatinization on the Xi through trimethylation of H3K27 ^11, 13, 35^. Earlier reports provided evidence that loss of Xist during XCI maintenance leads to alterations in H3K27me3 distribution across the Xi ^27, 36–39^. We hypothesized that reduced H3K27me3 on Xi may underlie transcriptional upregulation of X-linked genes in Xist-deficient HSPCs. To investigate, we first performed H3K27me3 immunostaining combined with X-paint DNA FISH on metaphase chromosomes of Lin- cells isolated from bone marrow of WT and Xist XistΔ/+ mice. As expected, we detected enrichment of H3K27me3 on the Xi in WT cells. In contrast, ∼45% of cells in XistΔ/+ Lin- cells presented complete loss of H3K27me3 on the Xi (Figure 4A). To investigate genome-wide changes in H3K27me3 distribution, we performed H3K27me3 ChIP-Seq in bone marrow Lin- cells isolated from WT and XistΔ/Δ female mice (Figure 4B). In Xist mutant cells, H3K27me3 levels were significantly reduced around the transcription start sites (TSS) and across gene bodies of X-linked genes (Figure 4B-C). Notably, reduction of H3K27me3 signal at X-linked DEGs was independent of their transcriptional state (Figure 4C). Moreover, we found that some XCI escape genes (e.g. Eif2s3x, Kdm6a and Kdm5c) were void of H3K27me3 in both WT and Xist deficient Lin- cells (Figure 4E). Based on these results, we concluded that loss of H3K27me3 likely contributes to the transcriptional changes across the X chromosome, but is not exclusively correlated with transcriptional upregulation of X-linked genes in Xist deficient Lin- cells.

**Figure 4:**
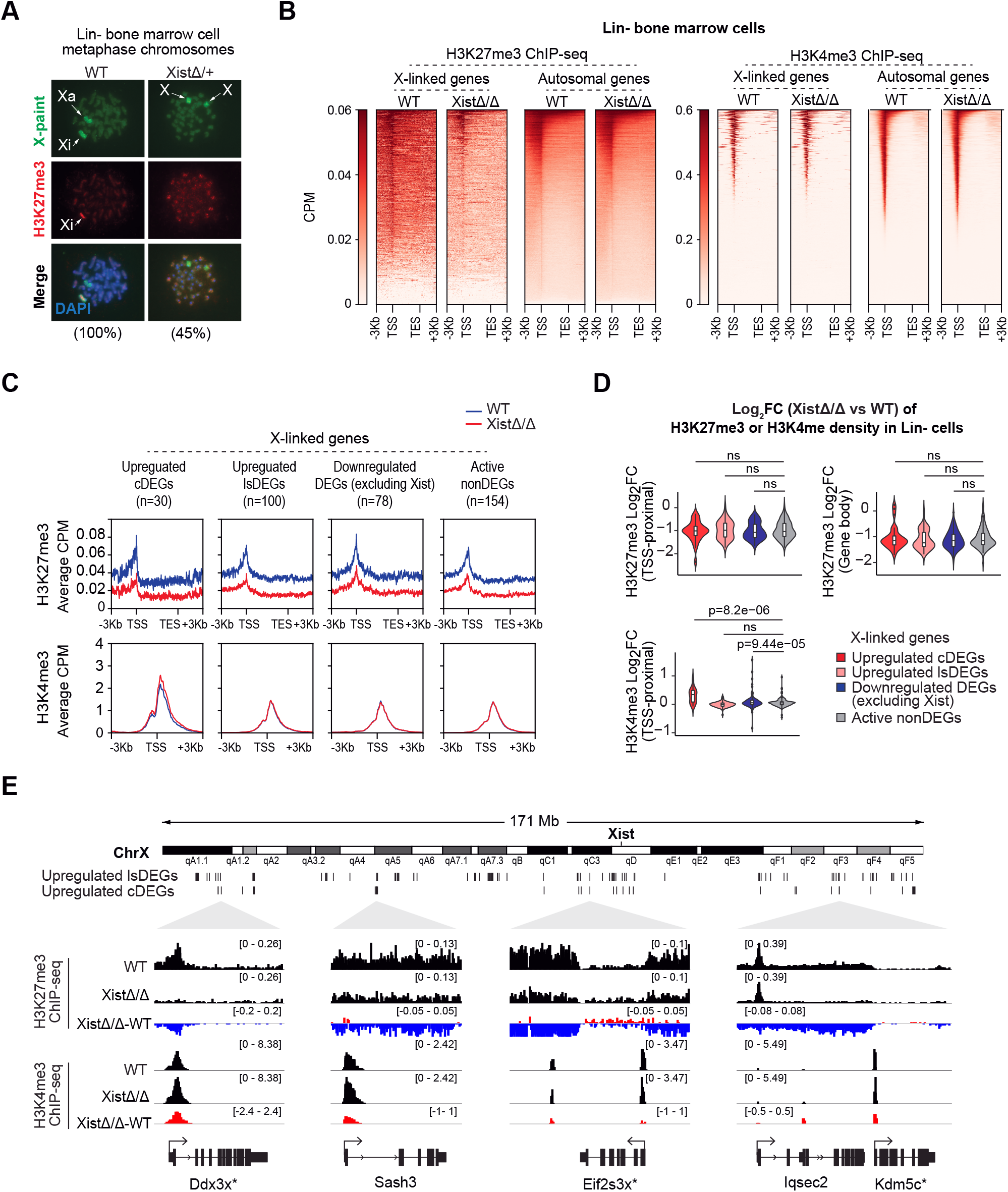
Xist deletion alters H3K27me3 and H3K4me3 distribution on X chromosome. **(A)** Representative images of sequential immunofluorescence (IF) staining of H3K27me3 (red) and DNA FISH of X-paint (green) on metaphase chromosomes of Lin- bone marrow cells isolated from 3- to 4- month-old WT and XistΔ/+ female mice. Xist deletion leads to loss of H3K27me3 signal on the Xi. The percentage of metaphase chromosomes with similar staining results are indicated. **(B)** Heatmaps of ChIP-seq identified H3K27me3 and H3K4me3 occupancy across +/- 3Kb of all X-linked and autosomal genes in Lin- cells isolated from 3-month-old WT and XistΔ/Δ female mice. Reduced H3K27me3 signal across X-linked genes was detected in Xist deficient cells. CPM, counts per million. **(C)** Metagene profiles show distribution of average H3K27me3 or H3K4me3 occupancy across +/- 3Kb of genes or +/- 3Kb of TSS of X-linked upregulated cDEGs and lsDEGs, downregulated DEGs, and active nonDEGs in Lin- bone marrow cells isolated from 3-month-old WT and XistΔ/Δ female mice. Number of genes in each group were indicated. **(D)** Violin and box plots show log2FC of H3K27me3 and H3K4me3 density at TSS-proximal sites (TSS +/- 3Kb) and H3K27me3 density within gene bodies of upregulated X-linked cDEGs and lsDEGs, downregulated DEGs (excluding Xist), and active nonDEGs in XistΔ/Δ cells in comparison to WT Lin- cells. Log2FC of H3K4me3 signal of upregulated X-linked cDEGs is significantly higher than X-linked active nonDEGs in Xist deficient cells. P-values from Student’s *t*-test were indicated; p ≥ 0.05 was defined as non-significant (ns). **(E)** Representative genome tracks show distribution of H3K27me3 and H3K4me3 ChIP-seq signal across five upregulated X-linked cDEGs in WT and XistΔ/Δ Lin- cells. The difference between two genotypes is represented by subtracting ChIP-seq signal of WT cells from ChIP-seq signal of XistΔ/Δ cells. Genes that have been reported as mouse XCI escape genes by Berletch et al. were marked by an asterisk (*).

### Transcriptionally upregulated X-linked genes associate with increased H3K4me3 density and chromatin accessibility in Xist deficient cells

H3K4me3 is a histone modification that associates with transcriptionally active promoters ^40, 41^. In contrast to drastic reduction in H3K27me3 distribution, we did not detect a significant overall change in H3K4me3 distribution across majority of the X-linked or autosomal genes in Xist deficient Lin- cells (Figure 4B-C). Instead, we detected a slight yet significant increase in H3K4me3 density around TSS of upregulated X-linked cDEGs (p=8.2e-06) in comparison to actively transcribed X-linked genes that do not show any change in expression (active nonDEGs) (Figure 4D and 4E). Recent studies revealed that alternations in H3K4me3 deposition around TSS associate with changes in transcriptional memory, stability, or termination, rather than directly affecting transcription ^42, 43^. Thus, we propose that increased H3K4me3 is likely an outcome of the active transcriptional state of the X-linked cDEGs. This finding suggests that additional factors are likely required to selectively trigger upregulation of X-linked genes in Xist deficient cells.

To determine potential factors that might contribute to the transcriptional regulation of X-linked genes, we first mapped changes in open chromatin regions by performing ATAC-seq ^44^ in WT and XistΔ/Δ LSK+ and LSK- cells (Figure S3A). As expected, we found that in WT cells the overall chromatin accessibility around TSS of X-linked genes is lower in comparison to the autosomal genes due to the facultative heterochromatin state of one of the two X chromosomes (Figure S3A). We next examined distribution of ATAC peaks around TSS-proximal (TSS +/- 3Kb) and TSS-distal sites (sites located beyond TSS +/- 3Kb) on X chromosomes and autosomes in LSK+ and LSK- cells (Figure S3B). We found that Xist loss did not result in a significant change in the number of TSS-proximal ATAC peaks, but resulted in a slight increase in the number of TSS-distal peaks on autosomes and X chromosomes in LSK+ and LSK- cells (Figure 5A). Next, we quantified the intensity of each consensus ATAC peak on X chromosomes (LSK+: 1563 peaks; LSK-: 1680 peaks) and autosomes (LSK+: 65055 peaks; LSK-: 7033 peaks) and defined differential ATAC peaks as the ones that show consistent increase or decrease in intensity in both biological replicates (Figure 5A and Table S2). We found higher percentage of increased chromatin accessibility sites compared to those that show decreased chromatin accessibility at both TSS-proximal and -distal sites in LSK+ and LSK- cell genomes (Figure 5B and S3B). In addition, LSK+ cells showed more increased chromatin accessibility sites compared to LSK- cells (Figure 5B and S3B). Lastly, by comparing changes in chromatin accessibility across X chromosomes and autosomes, we found that the percentage of increased chromatin accessibility sites on X chromosome is higher than those on autosomes in both LSK+ and LSK- cells (Figure 5B and S3B). These findings suggested that X chromosome is more susceptible to change compared to the autosomes, and such change influences X chromosome chromatin structure more robustly in LSK+ cells in comparison to LSK- cells.

**Figure 5:**
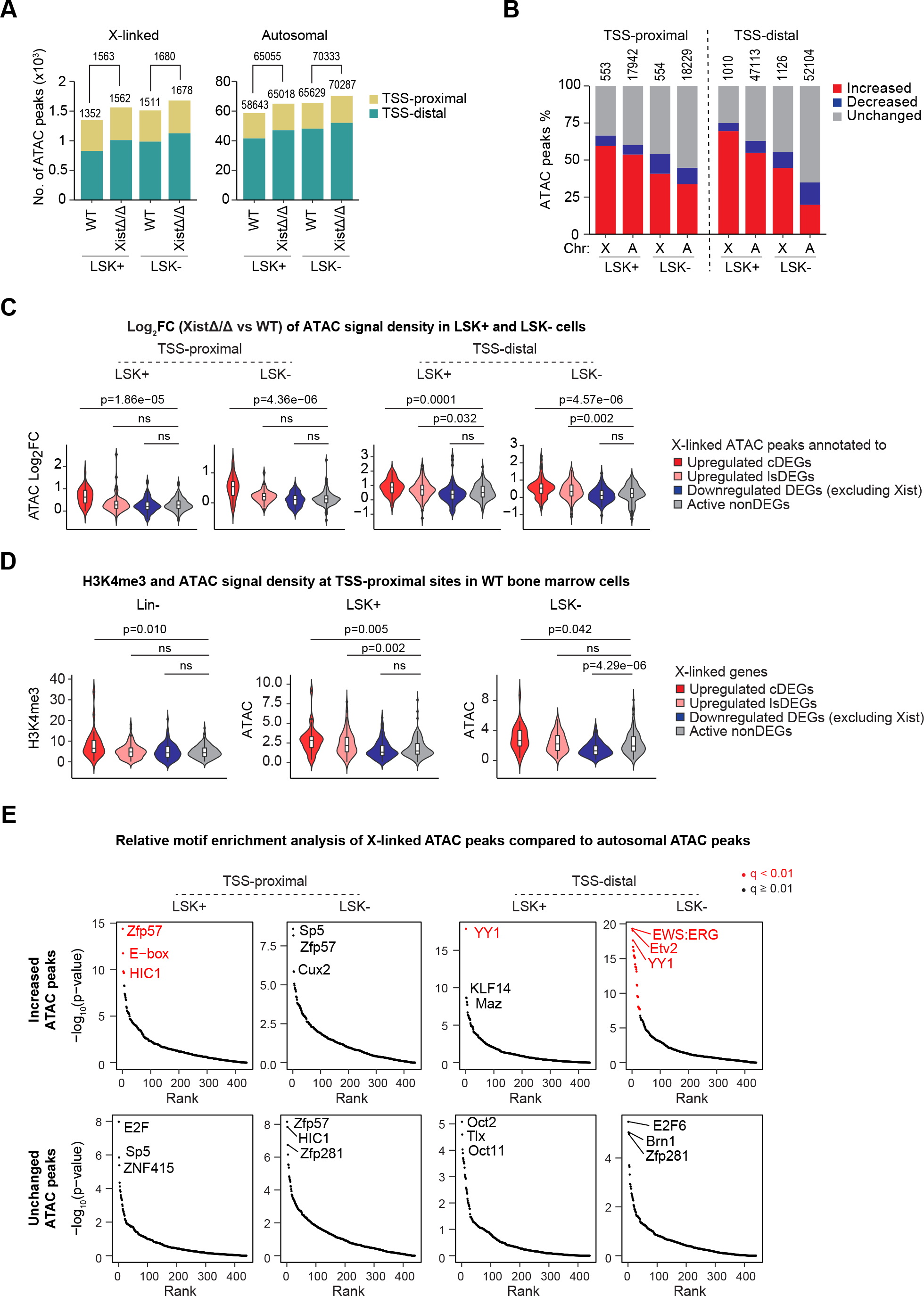
**Increased chromatin accessibility across X chromosome in Xist deficient cells**. **(A)** Bar graphs show number of ATAC peaks located at proximal- (TSS +/- 3Kb) and distal-TSS (peaks map beyond +/- 3Kb from TSS) regions on the X chromosome and autosomes in LSK+ and LSK- cells from 4.5-month-old WT and XistΔ/Δ female mice. Xist loss did not result in a significant change in the number of TSS-proximal ATAC peaks, but caused a slight increase in the number of TSS-distal peaks on autosomes and X chromosomes in LSK+ and LSK- cells. ATAC peak numbers are as indicated above corresponding bar graphs. **(B)** Bar graphs show percentage of increased, decreased, and unchanged ATAC peaks at proximal- and distal-TSS on X chromosome (X) and autosomes (A) in LSK+ and LSK- cells from 4.5-month-old WT and XistΔ/Δ female mice. The percentage of increased ATAC peaks on X chromosome is higher than that on autosomes in both LSK+ and LSK- cells. ATAC peak numbers are as indicated above corresponding bar graphs. **(C)** Violin and box plots show log2FC in intensity of TSS-proximal and TSS-distal ATAC peaks annotated to different types of X-linked DEGs and nonDEGs in LSK+ and LSK- cells from XistΔ/Δ cells in comparison to WT Lin- cells. Log2FC of ATAC-peak signal intensity annotated to upregulated cDEGs is significantly higher at TSS-proximal and -distal sites in Xist deficient LSK+ and LSK- cells. In contrast, upregulated lsDEGs exhibit significant increase in chromatin accessibility only at TSS-distal sites in LSK+ and LSK- cells. P-values from Student’s *t*-test were indicated; p ≥ 0.05 was defined as non-significant (ns). **(D)** Violin and box plots compare TSS-proximal H3K4me3 signal or ATAC signal of different types of DEGs and nonDEGs in WT cells. Upregulated cDEGs exhibited significantly higher H3K4me3 and ATAC signal compared to active nonDEGs on X chromosome. Upregulated lsDEGs exhibited significantly higher ATAC signal than active nonDEGs on X chromosome in LSK+ cell. P-values from Student’s *t*-test were indicated; p ≥ 0.05 was defined as non-significant (ns). **(E)** Relative motif enrichment analyses at increased or unchanged ATAC peaks located at X- linked proximal- and distal-TSS sites compared to autosomal sites. Motifs that are significantly enriched in X-linked increased ATAC peaks were highlighted in red (p<0.01 and q < 0.01). The names of top 3 enriched motifs were indicated in each plot. YY1 was detected as one of the candidate TFs.

To investigate the relationship between changes in chromatin accessibility and transcription of X-linked genes, we compared ATAC peak intensities across proximal- and distal-TSS of DEGs. In this analysis, we annotated each ATAC peak to a single gene based on proximal genetic distance (see Methods for details) (Table S2). We found that upregulated X-linked cDEGs showed significant increase in chromatin accessibility in Xist deficient LSK+ (p=1.86e-05) and LSK- (p=4.36e-06) cells compared to active nonDEGs at TSS-proximal regions (Figure 5C). Increase in chromatin accessibility was more widespread at TSS-distal sites whereby both upregulated cDEGs and lsDEGs associated with increased ATAC peaks in Xist deficient LSK+ and LSK- cells (Figure 5C). Furthermore, we found that transcriptionally upregulated X-linked genes were already enriched for H3K4me3 and ATAC signal compared to other X-linked genes in WT cells (Figure 5D). These data suggested that transcriptionally active and open chromatin sites on the X chromosome may serve as hot spots which could undergo alterations in chromatin structure in Xist deficient cells. Such changes could influence binding of transcription and/or chromatin regulatory factors.

### YY1 as a candidate transcription factor that associates with increased chromatin accessibility on X chromosome in Xist deficient cells

To identify candidate factors that might influence Xist-mediated regulation of transcription and chromatin accessibility on X chromosome, we performed HOMER motif enrichment analysis using increased ATAC peaks at TSS-proximal (LSK+: n=329; LSK-: n=226) and TSS-distal (LSK+: n=702; LSK-: n=502) regions of the X chromosome (Figure S3B and Table S2). In this analysis, increased ATAC peaks at autosomal TSS-proximal (LSK+: n=9643; LSK-: n=6137) and TSS-distal (LSK+: n=25893; LSK-: n=10337) regions were used as background control (Figure S3B and Table S2). The results showed that binding motifs of six factors, including Zfp57, E-box, HIC1, EWS:ERG, Etv2, and YY1, were significantly (p<0.01, q<0.01) enriched in X-linked increased ATAC peak sites at TSS-proximal or TSS-distal regions (Figure 5E and Table S2). Among these factors, YY1 stood out as a top candidate because its binding motif was significantly enriched at TSS-distal regions that associate with increased ATAC peaks on the X chromosome in both LSK+ and LSK- cells. Such enrichment was not present in the control motif analysis which was based on unchanged ATAC peaks (Figure 5E).

### Transcriptional upregulation of X-linked genes is associated with increased YY1 binding in Xist deficient cells

YY1 is an ubiquitous transcription factor which can influence transcriptional activation and repression ^45^. YY1 also acts as a bivalent protein that can bind with both DNA and RNA ^46^, and through such function it has been shown to be critical for nucleation of Xist onto the future Xi during the initiation of XCI ^12^. Beyond its direct role in transcription, YY1 has been shown to impact higher order chromatin organization by mediating DNA looping between enhancers and promoters ^47^. Based on these functional characteristics of YY1, we hypothesized that YY1 and Xist may co-regulate X-linked gene expression during XCI maintenance. To determine changes in YY1 binding in Xist deficient cells, we mapped genome-wide YY1 binding sites by ChIP-Seq in WT and XistΔ/Δ Lin- cells (Figure S4A and Table S3). We identified 346 X-linked and 13185 autosomal consensus YY1 binding sites in WT cells (Figure 6A and Table S3). We then determined differential YY1 peaks as those that exhibit consistent increase or decrease in YY1 binding intensity in both biological replicates of Xist deficient Lin- cells compared to their WT counterparts (Table S3). We found that, at the TSS proximal regions, Xist loss led to higher percentage of increased YY1 peaks (57%; 65/115) compared to decreased YY1 peaks (13%; 15/115) on the X chromosome but not on the autosomes (Figure 6B and S4B). This finding was in accordance with our motif analysis that showed enriched YY1 motif association with increased ATAC peaks in Xist deficient cells (Figure 5D). At the TSS-proximal regions, the percentages of increased and decreased YY1 peaks were comparable between the X chromosome and autosomes (Figure 6B). Nevertheless, metagene profiles showed that the change in the intensity of YY1 binding at increased peaks was larger than the change in YY1 binding intensity at decreased peaks on the X chromosome (Figure S4B). Based on these findings, we concluded that Xist deletion results in increased YY1 binding within both TSS-proximal and -distal sites across the X chromosome.

**Figure 6:**
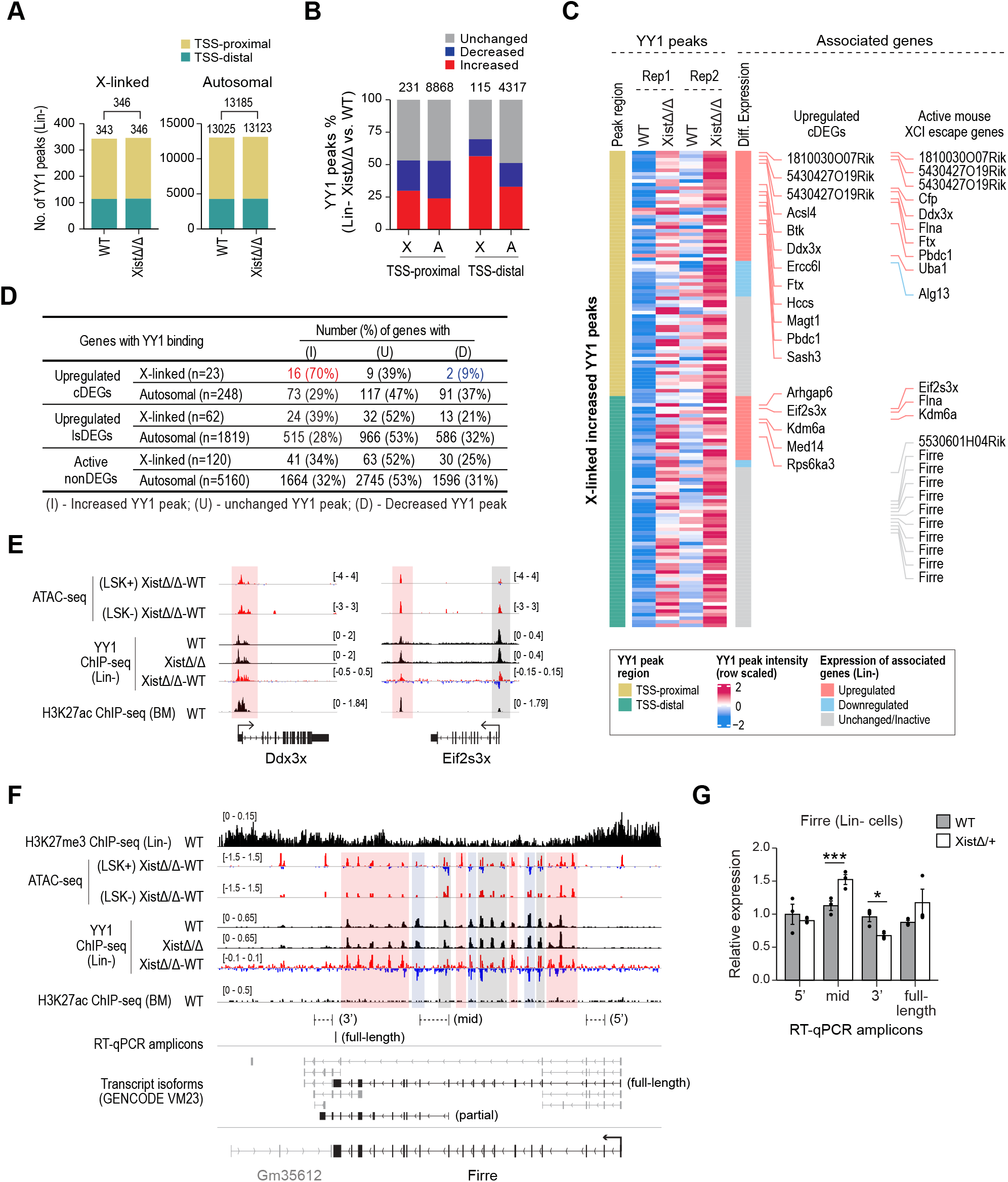
Transcriptional upregulation of X-linked genes is partially associated with increased YY1 binding in Xist deficient cells. **(A)** Bar graphs show comparable number of YY1 peaks at proximal- and distal-TSS on the X chromosome and autosomes in Lin- cells from 3-month-old XistΔ/Δ in comparison to WT female mice. Numbers of YY1 peaks are as shown. **(B)** Bar graphs show higher percentage of increased YY1 peaks at both TSS- distal sites on the X chromosome compared to autosomes in Lin- cells from 3-month-old XistΔ/Δ in comparison to WT female mice. Numbers of YY1 peaks are as shown. **(C)** Heatmap shows correlation between X-linked increased YY1 peaks and expression of associated genes at proximal- and distal-TSS regions. Names of upregulated X-linked cDEGs and active mouse XCI escape genes reported by Berletch et al. were indicated. **(D)** Table shows number and percentage of genes associated with increased, decreased, and unchanged YY1 peaks. Upregulated X-linked cDEGs show strong correlation with increased YY1 binding. **(E)** YY1, H3K27me3, H3K27ac ChIP-seq, and ATAC-seq tracks are shown for two upregulated X-linked cDEGs, Ddx3x and Eif2s3x. Increased (red) and unchanged (grey) YY1 peak regions were highlighted. **(F)** YY1, H3K27me3, H3K27ac ChIP-seq, and ATAC-seq tracks are shown for Firre locus. Increased (red), decreased (blue) and unchanged (grey) YY1 peak regions are highlighted. Splice variants of Firre gene and real time RT-PCR amplicon positions within the Firre locus are shown. **(G)** Real time RT-PCR showing relative Firre isoform mRNA levels in XistΔ/+ and WT bone marrow Lin- cells. GAPDH was used to normalize mRNA levels. Partial Firre transcript (see Figure 6F) shows upregulation in Xist deficient cells.

To investigate how differential YY1 binding is correlated with transcriptional changes in Xist-deficient cells, we annotated each YY1 peak to a single gene based on proximity in genetic distance (see Methods for details) (Figure 6C, S4C and Table S3). To examine TSS-distal YY1 binding sites in more detail, we also mapped potential enhancer sites by utilizing previously published Histone 3 Lysine 27 Acetylation (H3K27ac) ChIP-Seq in mouse bone marrow cells from the ENCODE portal ^48^ (https://www.encodeproject.org/) with the identifier ENCSR000CCL. Among the 23 upregulated X-linked cDEGs associated with YY1 binding, 16 genes (70%) exhibited increased YY1 peaks at either TSS-proximal or TSS-distal regions in Lin- cells (Figure 6C, 6D and 6E). Representative tracks are shown for two cDEGs, Ddx3x and Eif2s3x, in Figure 6E. We found that only 2 (9%) and 9 (39%) of the 23 upregulated X-linked cDEGs with YY1 binding associated with decreased or unchanged YY1 peaks, respectively (Figure 6D and S4C). Representative tracks for these genes, Nono and Utp14, are shown in Figure S4D. In contrast to the YY1 phenotype detected for upregulated X-linked cDEGs, we did not detect high percentages of genes that exhibit increased YY1 binding among upregulated autosomal cDEGs or lsDEGs (Figure 6D). These data indicated that there is a strong positive correlation between increased YY1 binding and transcriptional upregulation for a subset of upregulated X-linked cDEG in Xist- deficient cells.

We found that more than half (9/16) of the upregulated X-linked cDEGs that exhibited increased YY1 binding in Xist deficient cells (Figure 6C) have been previously characterized as XCI escape genes ^34^. Based on this finding, we investigated how Xist deletion affects YY1 binding and transcription of reported escape genes. In total, we detected 14 actively transcribed mouse XCI escape genes that associate with increased YY1 binding in our data set. Approximately 79% (11/14) of these genes showed transcriptional upregulation in Xist deficient Lin- cells (Figure 6C). In contrast, only 3 out of 12 (25%) XCI escape genes that are not associated with increased YY1 peaks (excluding Xist) showed transcriptional upregulation in Xist deficient Lin- cells (Figure S4C). We also noted that one of the XCI escape genes, Firre, ^49^ which fell under the group of genes that showed ‘unchanged transcription’ in Xist deficient cells (Table S1), exhibited multiple YY1 binding sites across its gene body which correlated with open chromatin regions (Figure 6F). This is unique from other X-linked genes that normally have one or two YY1 binding sites (Figure 6C, 6F and S4C). Firre encodes a lncRNA that have multiple splice isoforms ^49^ (Figure 6F). We found that several of the increased YY1 binding sites across the Firre gene-body were positioned to the vicinity of TSSs of different Firre splice isoforms (Figure 6F). Based on this finding, we hypothesized that changes in YY1 binding within the Firre gene body might associate with differential expression of a specific Firre isoform(s). To test this hypothesis, we performed real time RT-PCR and we found that Xist loss leads to upregulation of mid-region Firre isoforms and downregulation of the 5’ Firre isoforms (Figure 6G). The primers we used to detect mid-region isoforms can detect two different isoforms (full-length and partial) (Figure 6F-G). Using a second set of primers designed to uniquely capture full length Firre isoform, we found that this isoform was not differentially expressed in Xist-deficient Lin- cells (Figure 6G). This data indicated that it was the partial Firre isoform that is upregulated in Xist deficient cells.

Collectively, our results revealed that positive correlation between increased YY1 binding and transcriptional upregulation detected in Xist deficient cells is a unique feature of a large proportion of upregulated X-linked cDEGs, especially those that escape XCI. In our working model, we propose that Xist and YY1 cooperate to control expression of a subset of X-linked genes. During this cooperation, YY1 function in transcriptional activation of X-linked genes is limited in the presence of Xist, and such limitation would be released upon Xist depletion. These processes likely underlie the mechanism that governs transcriptional upregulation of X-linked genes in Xist deficient cells (Figure 7). Future studies are necessary to dissect the molecular basis of how Xist influences allelic YY1 activity and transcription on the X chromosomes.

**Figure 7:**
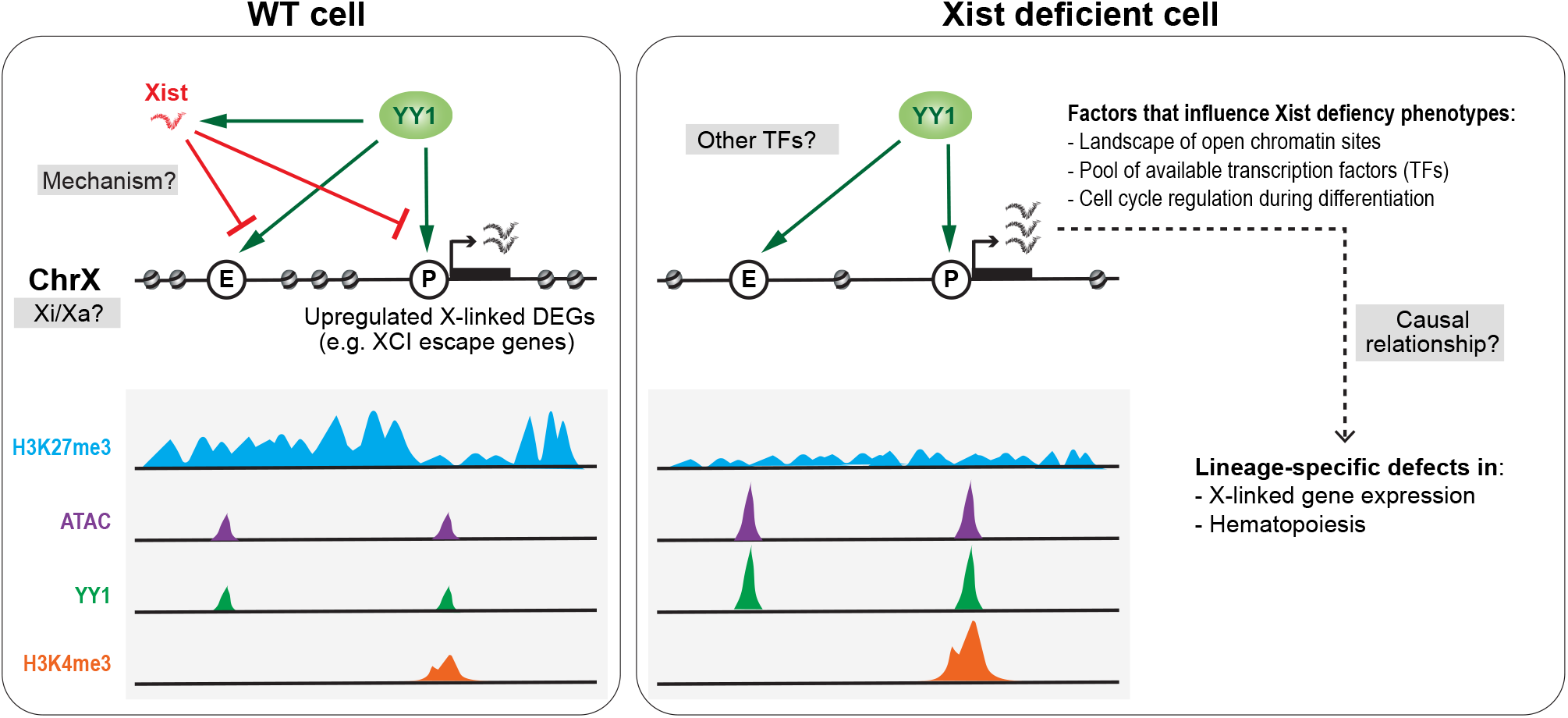
Working model on the regulatory role of Xist during XCI maintenance in hematopoietic cells. In our working model, we propose that Xist and YY1 cooperate to control expression of a subset of X-linked genes including XCI escape genes. Such cooperation involves limited transcription of X-linked genes through a YY1-dependent mechanism in the presence of Xist. Under conditions where Xist expression is altered, expression of this subset of genes is upregulated. Given that several of these genes have been implicated in immune response or regulation of hematopoiesis, it is likely that transcriptional upregulation of these genes is causal to cellular and pathological defects in Xist deficient mice. The severity of functional defects could be influenced by several factors including the landscape of open chromatin sites, pool of available TFs and regulation of cell cycle during differentiation of different cell lineages. Future studies addressing molecular basis of these processes and how they contribute to the pathologies that associate with Xist deficiency are necessary.

## DISCUSSION

XCI is a critical dosage compensation phenomenon that is essential for female development (Yang and Yildirim, 2017). Recent analyses of cancer genomes have shown that the X chromosome harbors approximately three-times more genes with potential to act as tumor suppressors than other chromosomes (Davoli et al., 2013), and the Xi accumulates two-to four-times more mutations than any other chromosome in cancer genomes (Jager et al., 2013). Thus, the Xi presents a critical target for treatment of X-linked disorders and XCI associated cancers and understanding molecular mechanisms impacting XCI maintenance would inform novel ways to transcriptionally reactivate the X-linked wild-type disease alleles for therapy.

### Cycling hematopoietic progenitors are sensitive to Xist loss during XCI maintenance

Initiation of XCI relies on Xist which is continuously expressed during the XCI maintenance. However, our understanding on the functional relevance of Xist during the maintenance state has been limited. This is because Xist function has been mainly investigated in cell culture. In this study, we analyzed the role of Xist in regulation of hematopoiesis, cell cycle and gene expression using the HSC-specific Xist knockout mouse model ^25^. In line with our previous findings, we detected expansion of LSK+ cells. Spefically, we found that at early disease stages, Xist deletion resulted in over-proliferation of pluripotent hematopoietic progenitors (HPC1/2), but not of HSCs which remain quiescent majority of the time. Our analyses revealed that the HPC1/2 phenotype is likely due to the additive systematic hematopoietic stress, especially at later disease stages. Nevertheless, we cannot exclude the possibility that Xist deletion is causal to transcriptional upregulation of an X-linked gene(s) that triggers cell signaling events leading to increased cell cycle activity. Our study also revealed that differentiation of common lymphoid progenitors is facilitated while differentiation of myeloid progenitor cells is obstructed in Xist deficient mice. These results are in accordance with the variable differentiation and maturation abnormalities associated with lymphoid and myeloid lineages in Xist deficient mice ^25^. Recent studies revealed that differentiation of HSCs into myeloid progenitors is less dependent on cell division in comparison to differentiation of HSCs into lymphoid progenitors ^50^. Therefore, the difference in cell cycle stress during differentiation of lymphoid and myeloid progenitors might underlie the variability in their tolerance to Xist deletion.

### X-linked genes at active chromatin sites are susceptible to transcription upregulation in Xist deficient cells

One of the interesting findings of our study is that we determined a subset of X-linked genes, among which many have been reported as XCI escape genes, that exhibit consistent transcriptional upregulation in all three HSPC populations of Xist deficient female mice. We referred to these genes as upregulated X-linked cDEGs. Because the bone marrow cells isolated from our mouse model is a mixture of cells with either paternal or maternal X chromosome being silenced, we were unable to determine allele-specific expression of this subset of X-linked genes using our RNA-seq data. Instead, we examined these loci for H3K27me3 and H3K4me3 occupancy, and chromatin accessibility in WT HSPCs. We determined lack of H3K27me3 within gene body of several transcriptionally upregulated XCI escape genes (Figure 4E). We also showed that in general, upregulated X-linked cDEGs presented high H3K4me3 occupancy around TSS (Figure 5D). Furthermore, we found that upregulated X-linked cDEGs were associated with open chromatin sites at TSS-proximal and -distal regions in WT LSK+ and LSK- cells (Figure 5D). Based on these results, we concluded that X-linked genes located at active chromatin sites, such as the XCI escape genes, may serve as hot spots for chromatin structure changes and transcriptional upregulation upon Xist depletion.

In addition to the upregulated X-linked cDEGs, we determined hundreds of X-linked genes that are transcriptionally upregulated in one or two HSPC populations in Xist deficient female mice, a gene set which we referred to as upregulated X-linked lsDEGs. A small proportion of this gene set have also been reported to escape XCI (Table S1). Importantly, we found that upregulated lsDEGs in LSK+ cells include more escape genes in comparison to upregulated lsDEGs in LSK- cells (Table S1). In line with this finding, upregulated X-linked lsDEGs show significantly higher chromatin accessibility compared to active nonDEGs in WT LSK+ cells, but not in WT LSK- cells (Figure 5D). Based on these results, we concluded that differences in the chromatin structure of the X chromosome in different cell types (i.e. LSK+ and LSK- cells) likely underlies the variability in transcriptionally upregulated X-linked genes upon Xist depletion.

### Binding of specific TFs influences upregulated transcription of X-linked genes in Xist deficient cells

Defining mechanisms that regulate transcriptional upregulation of X-linked genes in Xist- deficient cells is critical in understanding the functional role of Xist during XCI maintenance. In this study, we explored the mechanism by investigating changes in histone modifications and chromatin accessibility, and determined how these changes associate with binding of specific TFs and/or influence X-linked gene transcription. We found that Xist deletion resulted in chromosome wide loss of H3K27me3 and increased chromatin accessibility at selected sites across the X chromosome. Importantly, we determined YY1 as a candidate TF that showed increased binding at sites with increased chromatin accessibility (Figure 6). YY1 is one of the TFs that has been shown to activate Xist expression and Xist localization ^12, 51–53^. Beyond these studies, our understanding on the role of YY1 during XCI maintenance is limited. Here, we provide new evidence showing that Xist loss leads to increased YY1 binding at the TSS-proximal (e.g. promoters) and TSS-distal (e.g. enhancers) regions of upregulated X-linked cDEGs, especially for those that have been reported as XCI escape genes (Figure 6E-F). Our working model proposes coordinated function of Xist and YY1 in regulation of X-linked gene expression during XCI maintenance (Figure 7). To study how Xist controls allelic YY1 binding, future studies that utilize a mouse model that inherits fully skewed XCI by artificial manipulation and carries enough SNPs to distinguish individual alleles of X chromosomes would be instrumental.

Beyond a large subset of upregulated X-linked cDEGs that show increased YY1 binding, many upregulated X-linked DEGs (e.g. Nono, Utp14a, Tmem29, and Tlr7) did not show increased YY1 binding at sites with increased chromatin accessibility in Xist deficient cells (Figure S4C and S4D). It is likely that transcription of these genes relies on Xist in combination with other TFs. In support of this argument, a recent study provided evidence that transcriptional activation of human TLR7 on Xi is dependent on a B-cell-specific TF, TRIM28, in a Xist deficient human lymphoblastic cell line ^54^. TLR7 has been reported to escape XCI specifically in human immune cells ^55, 56^. In our study, we have detected increased chromatin accessibility around Tlr7 promoter in LSK+ cells but not in LSK cells (Figure S4D), suggesting that increased TF binding at Tlr7 promoter upon Xist loss is cell type dependent. We propose that the pool of available TFs in specific cellular contexts might dictate the range of X-linked genes that are susceptible to transcriptional upregulation upon Xist deletion, and such differential changes may associate with cellular and pathological defects in Xist mutant mice.

### Transcriptionally upregulated XCI escape genes are involved in immune response and hematopoiesis

Based on the literature, several XCI escape genes that showed transcriptional upregulation in our Xist deficient mouse HSPCs have been implicated in immune response (e.g. Ddx3x, Usp9x, Uba1, Flna, Rnf128, Btk, Cxcr3, Tlr7) ^57–65^, hematopoiesis (e.g. Gprasp1) ^66^, and cell division (e.g. Kif4) ^67^ in humans. To define the causal role of ’key’ X-linked genes in hematologic malignancies including the MPN/MDS ^25^, future studies should be implemented whereby expression of specific genes is downregulated in Xist mutant cells or upregulated in WT cells (Figure 7).

In addition to genes with known immune-related functions in human, we found that Xist deletion resulted in upregulation of a specific transcript isoform of the X-linked Firre gene. Notably, we detected increased YY1 binding across the gene body of Firre locus. Firre has been characterized as a lncRNA ^49^ that is biallelically expressed in various mouse tissues ^34^ with highest expression in HSCs ^68^. Lack of H3K27me3 within the Firre locus in WT Lin- cells suggested that Firre might be an XCI escape gene in HSPCs (Figure 6F). Recent studies examining Firre- overexpressing and Firre-deficient mice showed that Firre impacts hematopoiesis, especially cells of lymphoid origin ^68^. Thus, Firre is one of the potential XCI escape genes differential expression of which might contribute to hematopoietic defects in Xist mutant female mice. Given that Firre is expressed in several tissues, it is plausible that coordinated activity of YY1 and Xist might be critical in regulating expression of cell type-specific Firre isoforms. Future studies involving characterization of the function of Firre isoforms in a cell type specific context as well as the mechanism of how YY1 regulates their transcription would be insightful.

In sum, our study revealed that Xist expression during XCI maintenance controls transcription of a subset X-linked genes including several escape genes through a YY1- dependent mechanism. Some of these XCI escape genes have essential functions in hematopoiesis and immune response and increase in their expressional level might be causally linked to the development of systematic hematopoietic defects in Xist deficient female mice. The severity of functional defects caused by Xist loss could be affected by factors including the landscape of open chromatin sites which might contain XCI escape genes, pool of available TFs, and cell cycle stress during differentiation of specific cell lineages (Figure 7). Overall, our findings provide insights for future studies in which gene regulatory mechanisms that govern variability in XCI escape gene expression and tissue-specific tolerance to Xist deletion could be investigated.

## MATERIALS AND METHODS

### Mice

*Xist*^2lox/2lox^ mice (129Sv4/Jae strain) and B6.Cg-*Commd10^Tg (Vav1-cre) A2Kio^*/J (Vav-iCre, Jax 008610) mice were obtained from the Mutant Mouse Resource and Research Center (MMRRC) at the University of North Carolina (UNC) and The Jackson Laboratory, respectively. Mice were screened by PCR for *Vav.Cre*, *Xist*^WT^, and *Xist*^2lox^ alleles using the following primer sets: *Vav.Cre* (522-1: 5′-CTT CTC CAC ACC AGC TGT GGA-3′, 522-2: 5′-GAC AGG CAG GGC CTT CTC TGAA-3′; 580 bp) and *Xist* (Xint3F: 5′-GGC CAG TTT CTG ACA CCC TA-3′, Xint3R: 5′-CAC TGG CAA GGT GAA TAG CA-3′; *Xist*^WT^ 200 bp, *Xist*^2lox^ 300 bp). Mouse husbandry and experiments were carried out as stipulated by the Duke University Institutional Animal Care and Use Committee (IACUC).

### Isolation of bone marrow cells and FACS

Bones (tibias and femurs) were dissected, and bone marrow (BM) was flushed with 1xPBS supplemented with 2% FBS (2% F/PBS) into a 50-ml tube (Falcon) using a 22.5 gauge needle. Cell suspension was pipetted up and down for several times and filtered through a 70μm cell strainer (Falcon, 352350) to obtain a single-cell suspension. Red blood cells in the cell suspension were lysed by ACK Lysing buffer (Gibco, A10492-01) according to the manufacturer’s instructions. All antibodies used for FACS staining were from BD Pharmingen unless noted. To label lineage-positive cells, the following antibody cocktail (1:200) containing biotinylated antibodies against CD11b (M1/70), Gr-1 (RB6-8C5), CD3ε (145-2C11), B220 (RA3-6B2), CD8*α* (53-6.7), CD4 (GK1.5), and TER-119 (TER-119) in combination with BV421-conjugated Streptavidin (BD, 5832259; 1:200) was used. To distinguish between LSK+ and LSK- cells, biotinylated lineage-specific antibody cocktail was used in combination with FITC-conjugated SCA-1 (D7; 1:100) and PE-conjugated c-KIT (2B8; 1:100) antibodies. To distinguish LSK+ subpopulations (HSC, MPP HPC1, HPC2), cells were further labelled with PE/Cy7-conjugated CD150 (TC15-12F12.2; Biolegend; 1:100) and APC/Cy7-conjugated CD48 (HM48-1; Biolegend; 1:100) antibodies. To distinguish LSK- subpopulations (MEP, GMP, CMP), cells were labelled with AF647-conjugated CD34 (RAM34; 1:100) and PE/Cy7-conjugated FcγR (2.4G2; 1:100) antibodies. To identify CLP, we stained cells with PE/Cy7-conjugated IL7R*α*(SB/199; 1:100). Labelled cells were filtered through a 70 μm cell strainer before flow cytometry analysis or sorting. 7-Amino-Actinomycin D (7-AAD, BD559925; 1:200) solution was used to label the nonviable cells. Flow cytometry analysis was conducted on BD FACSCanto II Cell Analyzer (BD) and data is analyzed using FlowJo software. Sorting was conducted on MoFlo Astrios Cell Sorter (BD) or MA900 Multi-Application Cell Sorter (Sony).

### Ki67 and BrdU cell cycle assays

For Ki67 cell cycle assay, bone marrow cells were labelled using biotin-conjugated lineage antibody cocktail in combination with PerCP-conjugated streptavidin (BD; 554064) and antibodies against SCA-1, c-KIT, CD150 and CD48 as described above. Labelled cells were fixed and permeabilized using Fixation/Permeabilization solution (BD, 554714) at 4°C for 20 min and washed with 1x BD Perm/Wash buffer. Cells were next incubated in 1x BD Perm/Wash buffer containing AF674-conjugated anti-Ki67 antibody (16A8; Biolegend; 1:100) at 4°C overnight. The next day, cells were washed with 1X BD Perm/Wash buffer and resuspend in 2% F/PBS. For BrdU cell cycle assay, the staining was performed as previously described ^33^. Briefly, 5-bromo-2’-deoxyuridine (BrdU) (Sigma, 10280879001) solution was peritoneally injected into live mouse at a concentration of 150mg/kg. One hour later, mouse was euthanatized, bone marrow cells were isolated and labelled using biotin-conjugated lineage antibody cocktail in combination with PerCP-conjugated streptavidin (BD, 554064; 1:200), and antibodies against SCA-1, c-KIT, CD150 and CD48 as described above. Labelled cells were fixed by incubating in Fixation/Permeabilization solution at 4°C for 20 min, permeabilized by Permeabilization Buffer Plus (BD, 561651) at 4°C for 10min, and re-fixed by Fixation/Permeabilization solution at 4°C for 5 min. Fixed and permeabilized cells were then incubated in 70U/ml DNase I (Roche, 4716728001) at 37°C for 1 hour before staining with APC-conjugated anti-BrdU antibody (Bu20a, Biolegend; 1:100) at room temperature (RT) for 20 min. Cells were washed with 1x BD Perm/Wash buffer between steps and resuspend in 2% F/PBS. For both Ki67 and BrdU cell cycle assays, DAPI solution (Thermo, 62248; 1:500) was added to cell suspension to stain DNA 5 min before flow cytometry analysis which was conducted on BD FACSCanto II Cell Analyzer (BD). Data is analyzed using FlowJo software.

### Preparation of metaphase chromosome spreads

Bone marrow cells from mice were labeled with a cocktail of biotin-conjugated lineage marker antibodies in combination with Streptavidin Particles Plus (BD, 557812) according to the manufacturer’s instructions. Lineage-positive cells were depleted using Cell Separation Magnet (BD, 552311) according to the manufacturer’s instructions to obtain lineage-negative (Lin-) bone marrow cells. To arrest cells at metaphase, ∼2.5x10^6^ Lin- cells were incubated in complete RPMI media (Gibco) containing 100ng/ml nocodazole at 37°C for 2 hours. Cells were then resuspended in 5 ml of 0.056M KCl and incubated at room temperature for 30 min. KCl treated cells were then resuspend in 1 ml of cold methanol: glacial acetic acid (3:1) solution and spun down at 1000 rpm for 10 min 4°C for fixation. The fixation step was repeated for three times, and every time 200 μl of solution was left when removing the supernatant. After the last fixation step, cells were resuspended in the remaining 200 μl of methanol: glacial acetic acid (3:1). The fixed metaphase cell suspension can be store at -20°C for at least one year. To prepare metaphase chromosome spreads, 20 μl of metaphase cell suspension was released onto a pre-chilled glass slide with the help of a pipette from 1.5-meter height, air dried at RT and used for immunofluorescence (IF) staining or DNA fluorescence in situ hybridization (FISH).

### RNA FISH, DNA FISH and immunostaining

Experiments were performed as previously described ^38^ with minor modifications. Xist RNA was detected using Xist plasmid (E6) that contains the intronic sequence between Xist exon 7 and 8. Single-stranded Xist riboprobe was generated by in vitro transcription using biotin-16-UTP (Roche, 11388908910) and MAXIscript SP6 kit (Invitrogen, AM1322). The RNA probe is diluted in hybridization buffer (50% Formamide, 2xSSC, 2 mg/ml BSA, 10% Dextran sulfate-500k) at a concentration of 10 ng/μl, denatured at 75°C for 10 min and pre-annealed at 42°C for 30-90 min before hybridization. To perform RNA FISH, 10-50x10^3^ sorted LSK+ cells were incubated in 100 μl RPMI media on each well of a 12-well glass slide (Fisher Scientific, 9991090) at 37°C for 30 min. The media was discarded gently, the attached cells were washed twice with ice-cold 1xPBS, and permeabilized in ice-cold CSK solution (10 mM PIPES/KOH (pH 8.6), 100 mM NaCl, 300 mM Sucrose, 1 mM MgCl_2_) containing 0.5% Triton X-100 for 10 min. Cells were then fixed in 4% paraformaldehyde (PFA) at RT for 10 min, dehydrated in ice-cold 70%, 90% and 100% EtOH sequentially for 2 min per step and then air dried at RT for 5 min. Approximately, 5-6 μl of denatured and pre-annealed Xist RNA probe was added onto each well of the slide. The slide was then covered with a clean coverslip, sealed with rubber cement, and incubated overnight at 37°C in a dark humid chamber. Slides were then washed 3 times using 2xSSC containing 50% Formamide at 45°C for 5 min followed by a single wash in 0.1xSSC at 45°C for 5 min. Slides are quickly rinsed with 1xPBS containing 0.2% Tween-20 at RT for 10 seconds. To detect biotin labeled Xist RNA probe, the slide was further processed for immunostaining using AlexaFluor647-conjugated streptavidin (Invitrogen). Briefly, the slide was washed in blocking solution (4xSSC, 1% BSA, 0.1% Tween-20) at 37°C for 10 min, and hybridized with AlexaFluor647-conjugated streptavidin (S21374; Invitrogen; 1:400 diluted in blocking solution) at 37°C for 40 min in a dark humid chamber. Cells were washed three times in 4xSSC, 0.1% Tween-20 solution at 45°C for 10 min each.

To perform H3K27me3 and H3Ser10P immunostaining of LSK+ cells following RNA FISH, the cells were incubated with immunostaining (IF) blocking solution (1xPBS, 1% BSA, 0.1% Tween-20) at RT for 1 hour. This was followed by hybridization step with primary antibodies including anti-H3Ser10P (3H10; Millipore; 1:200) and anti-H3K27me3 (C36B11; Cell Signaling; 1:200) diluted in IF blocking solution at RT for 1 hour. The slide was washed three times with 1xPBS, and then incubated with secondary antibodies including AlexaFluor555-conjugated goat anti-mouse IgG (A-21422; Invitrogen; 1:500) to detect H3Ser10P and AlexaFluor488-conjugated goat anti-rabbit IgG (A-11008; Invitrogen; 1:500) to detect H3K27me3 for 1 hour at RT, and then washed three times in1xPBS.

For metaphase chromosome DNA FISH, FITC labeled mouse X-paint (ASI) was used. To prepare the X-paint probe, 10 μl of X-paint solution was dried together with 10 μg of mouse Cot-1 DNA (Invitrogen, 18440-016) using Savant Integrated SpeedVac Concentrator and the pellet was resuspended using 50 μl (5x volume) of hybridization buffer. To perform sequential H3K27me3 immunostaining and X-paint DNA FISH using metaphase chromosome spreads, air dried metaphase spread slides were first blocked in IF blocking solution for 1 hour at 37°C, followed by incubation with anti-H3K27me2/me3 antibody (7B11; Active Motif; 1:200) for 3 hours at 37 °C. Slides were then washed three times in 1xPBS solution for 3 hours at 37°C and chromosome spreads were hybridized with Alexa Fluor555-conjugated goat anti-mouse IgG (1:500) for 1 hour at 37°C. After washing three times with 1XPBS, spreads were re-fixed with 4% PFA for 10 min at RT, denatured in 70% formamide/2xSSC buffer at 80°C for 10 min, dehydrated in ice-cold 70%, 90% and 100% EtOH sequentially for 2 min per step, and then air dried for 5 min. X-paint probe (10 μl) was denatured at 80°C for 10 min, pre-annealed at 37°C for 30-90 min and was added onto each slide. Probe solution was covered using a clean 18mmx50mm coverslip, sealed with rubber cement, and then incubated overnight at 37°C in a dark humid chamber. Slides were sequentially washed three times with 50%Formamide/2xSSC at 45°C for 5 min, three times in 2xSSC at 45°C for 5 min, and once in 0.1xSSC at 45°C for 5 min.

As a last step of FISH and IF protocols, air dried cells or chromosome spreads were mounted using mounting media containing DAPI (Vectashield). A cover slip was placed on the slide and sealed with nail polish before imaging. Images were acquired on a Leica Automated Upright Microscope (Leica, DM5500B) and analyzed using Leica Application Suite X (LAS X) and FiJi software.

### Total RNA extraction, reverse transcription and real-time PCR

Mouse Lin- bone marrow cells were collected by depleting lineage positive cells as described above and total RNA was isolated using TRIzol reagent (Invitrogen; 15596026) according to the manufacturer’s instructions. To isolate total RNA from LSK+ and LSK- cells, RNeasy Plus Micro kit (Qiagen, 74034) was utilized. For reverse transcription, cDNA was prepared using M-MLV Reverse Transcriptase (Invitrogen, 28025013) with random hexamers (Invitrogen, 48190011). Real-time PCR (qPCR) was performed using iTaq Universal SYBR Green Supermix (Bio-Rad, 172-5124) with the following primer sets. Firre 5’ transcript: (JR2-F: 5’- CAAATTCAAGCAGGCAAGGG-3’; JR2-R: 5-AGGTATGCTTCACCTCTCCT-3’); Firre mid transcript: (CD-F: 5’-TTCCTCATTCCCCTTCTCCT-3’; CD-R: 5’-ACCAGGTACCGTGAGCAATC- 3’) ^69^; Firre 3’ transcript: (TY-F: 5’-TGTCCATCCTTATCCAGGTGC-3’; TY-R: 5’-TGTGGGCACCCAAGTCATTA-3’); Firre full transcript: (JR4-F:5’- AACAGTGCCCATTTCAGTCC-3’; JR4-R: 5’- TTTTTCATGCAGGGTGATTG-3’) ^70^. Relative gene expression was calculated by the relative standard curve method. GAPDH expression was used to normalize data.

### RNA-Seq

Total RNA from WT and XistΔ/Δ mouse bone marrow Lin- cells (200x10^3^) was extracted using TRIzol reagent (Invitrogen), and RNA from LSK+ or LSK- cells (∼30-50x10^3^) was extracted using RNeasy Plus Micro Kit (Qiagen, 74004). RNA quality and concentration were assessed using a 2100 Bioanalyzer (Agilent Technologies) and Qubit 2.0 (Thermo Fisher Scientific), respectively. RNA-seq libraries of Lin- cells were prepared using Ovation RNA-Seq System Kit (NuGen, 0403-32 and 0348-32), and LSK+ or LSK- RNA-seq libraries were prepared using SMART-Seq v4 Ultra Low Input RNA Kit (Takara, 634888) and KAPA HyperPrep kit (KAPA, 07962347001). Before sequencing, fragment length distribution and library quality were first assessed on a Fragment Analyzer (Agilent). All libraries were pooled in equimolar ratio and sequenced. RNA-seq libraries were sequenced at 100 bp single-end (SE) on Illumina HiSeq 2500 for Lin- cells, 50 bp SE on Illumina HiSeq 4000 for LSK+ cells and 50 bp pair-end (PE) on Illumina NovaSeq 6000 for LSK- cells. Two biological replicates of RNA-seq experiments were conducted for each cell type and genotype.

### Chromatin immunoprecipitation sequencing (ChIP-seq)

ChIP experiments were performed as previously described with minor changes ^71^. Briefly, to perform H3K27me3 and H3K4me3 ChIP, mouse bone marrow Lin- cells (∼3x10^6^) were crosslinked with 1% formaldehyde for 10 min at RT and the reaction was stopped by adding glycine (final concentration, 125 mM) for 5 min at RT. Crosslinked cells were treated with hypotonic buffer (10 mM HEPES-NaOH, pH 7.9, 1.5 mM MgCl_2_, 10 mM KCl, 0.2 % NP-40, 1 mM DTT, 1 mM PMSF, 1x protease inhibitor cocktail (Sigma, P8340)) at 4°C for 10 min and collected in a tube. The cells were lysed in 150 μl of ChIP lysis buffer (50 mM Tris-HCl, pH 7.9, 10 mM EDTA, 1% SDS, 1 mM PMSF, protease inhibitor cocktail (Sigma, P8340)) on ice for 10 min, and sonicated for 35 cycles (30’’ ON + 30’’ OFF) using Diagenode Bioruptor to shear the chromatin to 200-1,000 bp fragments. The lysate was diluted with 9x volumes of ChIP dilution buffer (16.7 mM Tris-HCl, pH 7.9, 167 mM NaCl, 1.2 mM EDTA, 1.1% Triton X-100, 1 mM PMSF). The fragmented chromatin was pre-cleared using 15 μl of pre-washed Protein A/G agarose beads (Thermo Fisher, 20423) for 1 hour at 4°C. The pre-cleared chromatin fragments were equally separated into two aliquots. Each aliquot was incubated with either 2 μl of anti-H3K27me3 (mAbcam6002; Abcam) or 2 μl of anti-H3K4me3 (ab8580; Abcam) antibodies at 4°C overnight with rotation, followed by incubation with pre-washed Protein A/G agarose beads (Thermo Scientific, 20423) for 1-2 hours at 4°C. DNA-protein complexes attached to agarose beads were washed at 4°C with the following buffers: low salt wash buffer (20 mM Tris-HCl, pH 7.9, 100 mM NaCl, 2 mM EDTA, 0.1% SDS, 1% Triton X-100) for 10 min, high salt wash buffer (20 mM Tris-HCl, pH 7.9, 500 mM NaCl, 2 mM EDTA, 0.1% SDS, 1% Triton X-100) for 10 min, the LiCl buffer (10 mM Tris-HCl, pH 7.9, 250 mM LiCl, 1% NP-40, 1% deoxycholate acid, 1 mM EDTA) for 10 min, and twice with TE buffer for 5 min. DNA-protein complexes were eluted by incubating the beads in elution buffer (1% SDS, 0.1 M NaHCO_3_) for 15 min at RT. Reverse crosslinking was performed by incubating samples in 200 mM of NaCl at 65°C overnight followed by Proteinase K (20 μg/ml) treatment for 2 hrs at 55°C. DNA was purified using MiniElute PCR purification kit (QIAGEN, 28006).

To perform YY1 ChIP, bone marrow Lin- cells (∼1.5x10^6^) were crosslinked with 1% formaldehyde for 15 min at RT and the reaction was stopped by adding glycine (final concentration, 125 mM) for 5 min at RT. Crosslinked cells were treated with cell lysis buffer (10 mM Tris-HCl pH 7.5, 10 mM NaCl, 3 mM MgCl_2_, 0.5% NP-40, 1 mM DTT, 1 mM PMSF, 1x protease inhibitor cocktail (Sigma, P8340)) for 10 min at 4°C. To digest chromatin into <1,000 bp fragments, the pellet was washed twice with MNase reaction buffer (10 mM Tris-HCl pH 7.5, 10 mM NaCl, 3 mM MgCl_2_, 1 mM CaCl_2_, 0.2% NP-40, 1 mM DTT, 1 mM PMSF, 1x protease inhibitor cocktail), resuspended in 200 μl of MNase reaction buffer containing 2.5U of MNase (NEB, M0247) and incubated for 10 min at 37°C. MNase digestion was stopped by adding 20 μl of 0.5M EDTA into the reaction. Nuclear membrane was broken up by sonicating the lysate for 3 cycles (output level 6, 10’’ ON + 20’’ OFF) using Microson ultrasonic cell disruptor (Misonix). Lysate was spun at 21,000x*g* for 10 min at 4°C, and the supernatant containing fragmented chromatin was incubated with 2 μl of anti-YY1 antibody (D5D92Z, Cell Signaling, 46395) overnight at 4°C. DNA-protein complexes were pulled down with addition of 10 μl of pre-washed Protein G Magnetic Beads (Invitrogen, 10003D) and incubation for 1-2 hours at 4°C. DNA-protein complexes were eluted in elution buffer (1% SDS, 0.1 M NaHCO_3_) by incubating at 65°C for 40 min, and reverse crosslinked as described above.

All ChIP DNA samples were purified using MinElute PCR purification kit (Qiagen,28006) and quantified using the fluorometric quantitation Qubit 2.0 system (Thermo Fisher Scientific). ChIP-Seq libraries were prepared using KAPA HyperPrep kit (KAPA, 07962347001). Libraries were then pooled into equimolar concentration and sequenced on an Illumina HiSeq 4000 instrument (50 bp SE; H3K27me3, and H3K4me3 ChIP-Seq) and on an Illumina NovaSeq 6000 instrument (50 bp PE; YY1 ChIP-Seq). Two biological replicates of ChIP experiments were conducted for each cell type and genotype.

### Assay for Transposase Accessible Chromatin with high-throughput sequencing (ATAC-seq)

ATAC experiments were performed as previously described with minor modifications ^44^. Briefly, bone marrow LSK+ or LSK- cells (∼1x10^4^) were lysed in ice-cold lysis buffer (10mM Tris-HCl, pH 7.4, 10mM NaCl, 3mM MgCl_2_, 0.1% NP-40) for 5 min and then washed twice with 10mM Tris-HCl, pH 7.4. Following the last wash, 25 μl of extracted nuclei was mixed with 25 μl of 2XTD buffer (66 mM Tris-Ac pH 7.8, 132 mM KAc, 20 mM MgAc, 32% Dimethyl Formamide) containing 1 μl of Tn5 with loaded tagmentation adapters. Transposition reaction mixture was incubated using a thermocycler rotating at 500 rpm for 1 hour at 37°C. Tagmented DNA fragments were purified using MinElute PCR Cleanup Kit (Qiagen, 28206) and amplified for 15 PCR cycles using Q5 polymerase (NEB, M0491). PCR products were run on a gel, and fragments of ∼200-800 bp size were isolated and sequenced on MGI DNBseq-G400 or Illumina NovaSeq 6000 platform at 50bp PE. Two biological replicates of ATAC experiments were conducted for each cell type and genotype.

### RNA-seq data analysis

RNA-seq reads were trimmed using TrimGalore (v0.6.0, -q 15) ^72^ and then aligned to mm10 reference genome by TopHat (v2.1.1, --b2-very-sensitive --no-coverage-search) ^73^. Reads with low alignment quality were filtered out using Samtools (v1.10, -q 30) ^74^. Aligned reads for each gene in UCSC mm10 GTF file were counted by featureCounts (v1.6.3) ^75^ and normalized using DEseq2 (v1.30.1) ^76^. Transcript from at least 16900 (69%) genes were detected for each sample. Genes with FPKM > 1 in WT or XistΔ/Δ or both genotypes are defined as active genes. Active genes showing consistent increase or decrease in read counts in both biological replicates were determined as differentially expressed genes (DEGs), while the remaining active genes are determined as non-differentially expressed genes (nonDEGs). Gene Ontology (GO) analysis was conducted using clusterProfiler R package (v3.18.1) ^77^. Gene set enrichment analyses (GSEA) was conducted using the GSEA software (v4.0.3) with MSigDB ontology gene sets (v6.2) or customized signature gene sets of myeloid progenitors ^31, 78^. To obtain signature genes for myeloid progenitors, published RNA-seq data for CMPs, GMPs and MEPs (GSE60103) ^79^ was utilized. Genes transcription of which were significantly upregulated (p<0.05 and FPKM>1) in one myeloid progenitor population, while not actively transcribed (FPKM < 1) in the other two myeloid progenitor populations were determined as ’signature genes’ (See Figure S1C for heatmap). To compare Xist deletion-induced DEGs between tissues, RNA-seq data of other tissue-specific Xist- mutant mouse models were obtained from European Bioinformatics Institute’s repository under accession number E-MTAB-8161 (kidney, crypt and polyp) ^28^ and from GEO database under accession number GSE97077 (brain) ^27^. Two biological replicates of published RNA-seq results of each tissue-specific Xist-mutant mouse model were used for analyses using the same methods as our HSPC RNA-seq data.

### ChIP-seq data analysis

ChIP-seq reads were trimmed using TrimGalore and then aligned to mm10 reference genome by Bowtie2 (v2.3.5.1, --qc-filter --very-sensitive) ^80^. Reads with alignment quality lower than 30 or reads aligned were removed using Samtools (v1.10). For H3K27me3 and H3K4me3 ChIP-seq, at least 50x10^6^ high-quality aligned reads were obtained for each sample. For YY1 ChIP-seq, duplicated reads were removed by MarkDuplicates tool of Picard (v2.0.1; http://broadinstitute.github.io/picard/). At least 40x10^6^ high-quality aligned reads were obtained for each sample after removal of duplicates. ChIP-seq results of two biological replicates were merged to make bigwig files, metagene profiles and heatmaps using deepTools (v3.5.0) ^81^ with CPM normalization method. To map potential enhancer regions, H3K27ac ChIP-seq result in bone marrow cells (ENCSR000CCL) was obtained from ENCODE database ^48^ (https://www.encodeproject.org/) and bigwig files were generated in the same way as our ChIP-seq data.

For H3K27me3 and H3K4me3 ChIP-seq, aligned reads within TSS-proximal (TSS ± 3Kb) region for each gene in UCSC mm10 GTF file were counted by multicov program of bedtools (v2.25.0) ^82^. The density of H3K27me3 or H3K4me3 signals was calculated using the following formulas:

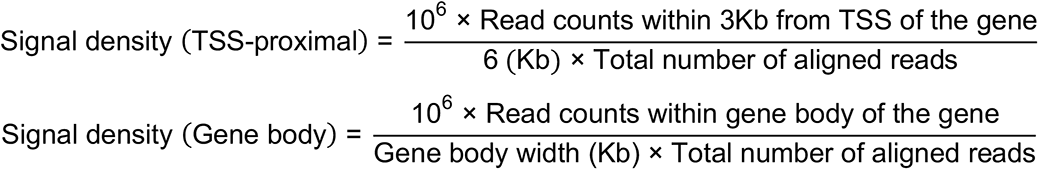

For YY1 ChIP-seq, peaks were called using findPeaks (-region -size 500 -minDist 500 -F 0 -L 4 -C 2) program in HOMER (v4.9.1) ^83^ and consensus YY1 peak sites from all samples were calculated by DiffBind R package (v3.0.13, minOverlap = 2; https://bioconductor.org/packages/DiffBind/). Consensus peaks called in no less than 50% of samples were considered as real YY1 peaks. To quantify intensity of each YY1 peak, reads within each YY1 peak site were counted (bUseSummarizeOverlaps = TRUE, summits = TRUE, filter = 0) and normalized (method = DBA_DESEQ2) using DiffBind. Intensity of each YY1 peak was calculated using the following formula:

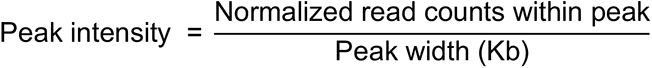

YY1 peaks that show consistent higher or lower intensity in Xist deficient cells compared to WT cells in both experimental replicates were defined as increased or decreased YY1 peaks, respectively. Each YY1 peak was annotated to one gene in EnsDb.Mmusculus.v79 annotation database (https://bioconductor.org/packages/EnsDb.Mmusculus.v79/) using ChIPpeakAnno R package (v3.24.1, select="bestOne") ^84^. YY1 peaks within TSS-proximal regions were first annotated to a gene with the closest TSS. Next, TSS-distal YY1 peaks were annotated to a gene with the closest gene body region.

### ATAC-seq data analysis

ATAC-seq reads were trimmed using TrimGalore and then aligned to mm10 reference genome by Bowtie2 (v2.3.5.1, --qc-filter --very-sensitive). Reads with alignment quality lower than 30 or reads aligned to mitochondrial DNA were removed using Samtools (v1.10). At least 20x10^6^ high-quality aligned reads were obtained for each sample after removing mitochondrial DNA. ATAC-seq bigwig files, metagene profiles and signal density within TSS-proximal regions were all processed in the same way as ChIP-seq. ATAC peaks were called using callpeak program (-q 0.01 -B -f BAMPE) in MACS2 (v2.2.7.1) ^85^ and consensus ATAC peak sites were calculated by DiffBind. Consensus ATAC peaks called in no less than 50% of samples of the same cell type were considered as real ATAC peaks. Intensity and annotated gene for each ATAC peak were obtained using the same method as YY1 peaks. ATAC peaks with consistent higher or lower intensity in Xist deficient cells compared to WT cells in both experimental replicates were defined as increased or decreased ATAC peaks. Motif analyses of sequences associated with differential ATAC peaks was conducted using findMotifsGenome program (-size given -len 8,10 -mis 2 -h - mask) of HOMER.

### Statistical analysis

Significance of the difference between control and knockdown cells for variables was analyzed with parametric Student’s *t*-test.

### Data availability

Raw and processed sequencing data have been deposited in the Gene Expression Omnibus (GEO) under accession number GSE184776.

## ACKNOWLEDGEMENTS

We are grateful to the members of the Yildirim lab for stimulating discussions and feedback. The authors thank Y. Jeon, S. Nechaev, M. Aladjem, and S. Kadota for critical reading and feedback on the manuscript. We thank Y. Diao Lab at the Department of Cell Biology at Duke School of Medicine for providing the tagmented Tn5 transposase. We thank N. Devos at the Duke Genome Sequencing Shared Resource for assistance in ChIP-, RNA-Seq library preparation and sequencing, L. Martinek and B Li at the Duke FACS Shared Resource for assistance in sorting of mouse bone marrow cells. This work was supported by the Whitehead Scholar Award (E.Y.).

## AUTHOR CONTRIBUTIONS

E.Y. conceptualized and supervised the study. T.Y., and E.Y. designed the experiments. T.Y. performed all aspects of experiments. T.Y. and J.O. analyzed sequencing data together with E.Y. T.Y. and E.Y. wrote the paper with input from J.O.

## COMPETING INTERESTS

The authors declare no competing interests.

## MATERIALS & CORRESPONDENCE

Further information and requests for resources and reagents should be directed to and will be fulfilled by the Lead Contact, Eda Yildirim.

## SUPPLEMENTAL INFORMATION

**Figure S1:**
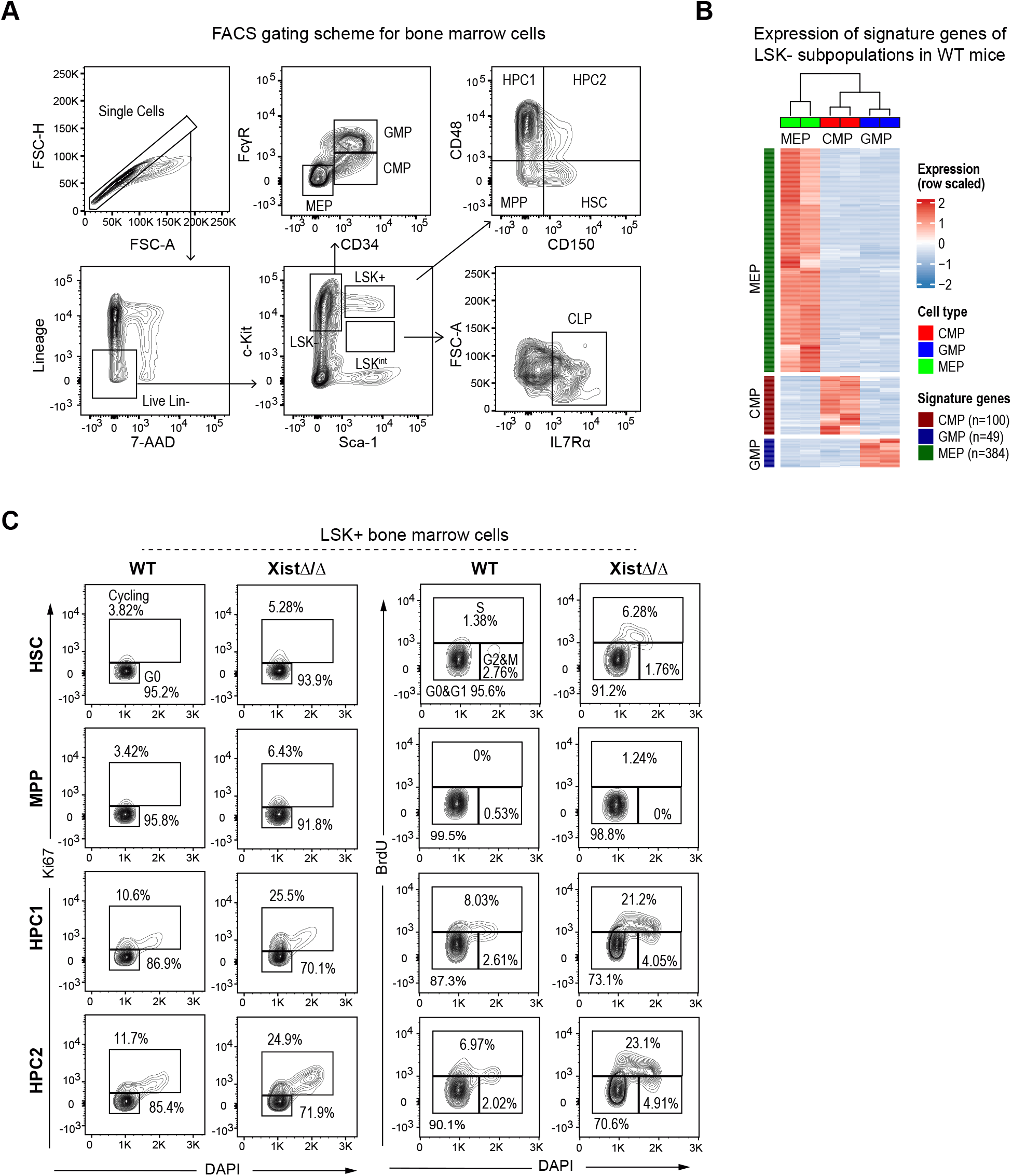
Gating schemes for identification of cell types and cell cycle stage of HSPCs, and expression of signature genes of LSK-cell subpopulations. Related to **Figure 1**, **2, and 3**. **(A)** Representative gating scheme for identification of LSK+ cells, LSK- cells, LSK+ subpopulations (HSC, MPP, HPC1/2), LSK- subpopulations (CMP, GMP and MEP) and lymphoid progenitors (CLP) in mouse bone marrow by FACS. **(B)** Heatmap shows gene expression of LSK- subpopulation cell signature genes in WT mice. **(C)** Representative FACS gating scheme for Ki67 and BrdU cell cycle analysis of LSK+ cell subpopulations from a pair of 3-month-old WT and XistΔ/Δ female mice.

**Figure S2:**
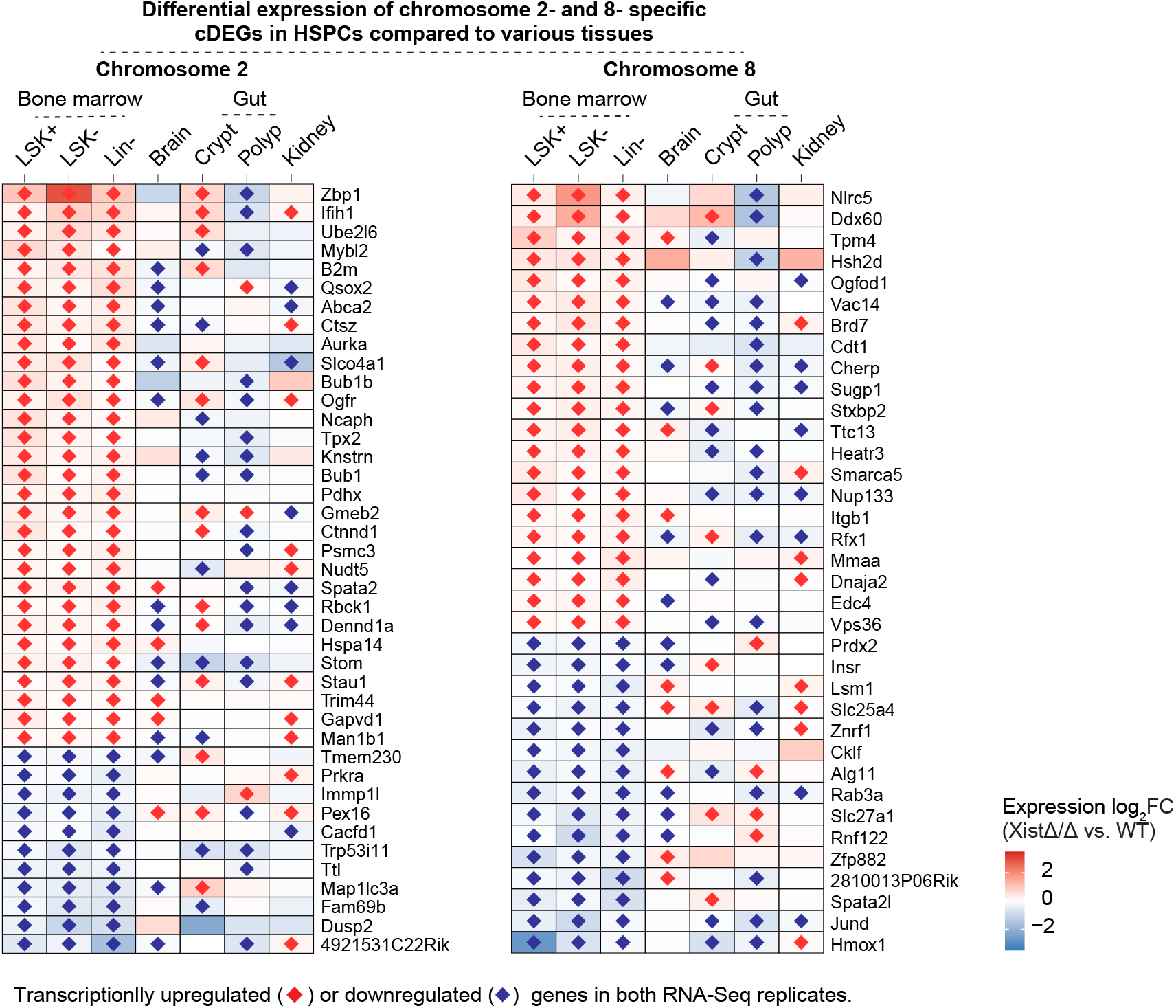
Differential transcriptional changes in chromosome 2 and 8-specific cDEGs in HSPCs compared to various tissues. Related to **Figure 3.** Heatmaps show expression of Log2FC of chromosome 2 and 8 linked cDEG in XistΔ/Δ and WT bone marrow cells (LSK+, LSK- and Lin- cells) in comparison to various tissues including brain ^27^, gut (crypt, polyp) and kidney ^28^.

**Figure S3:**
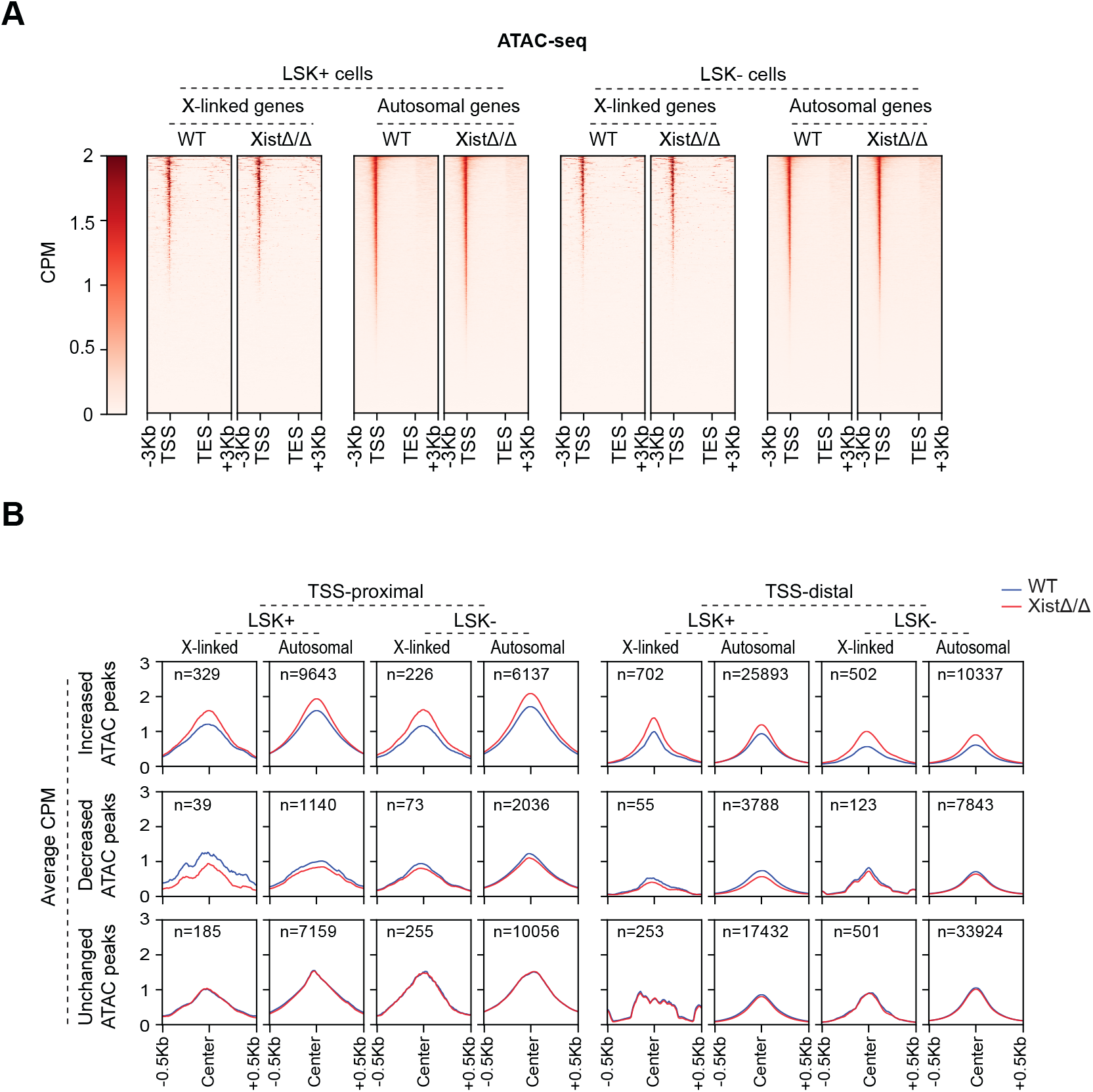
Analysis of chromatin accessibility by ATAC-seq in Xist deficient cells. Related to **Figure 5**. **(A)** Heatmaps of ATAC-seq signal +/- 3Kb from TSS of all X-linked and autosomal genes in LSK+ and LSK- cells from 3-month-old WT and XistΔ/Δ female mice. X-linked genes show lower chromatin accessibility in comparison to autosomal genes. CPM, counts per million. **(B)** Metagene profiles show distribution of average ATAC signal +/- 0.5Kb from the center of increased, decreased or unchanged ATAC peaks in WT and XistΔ/Δ LSK+ and LSK- cells.

**Figure S4:**
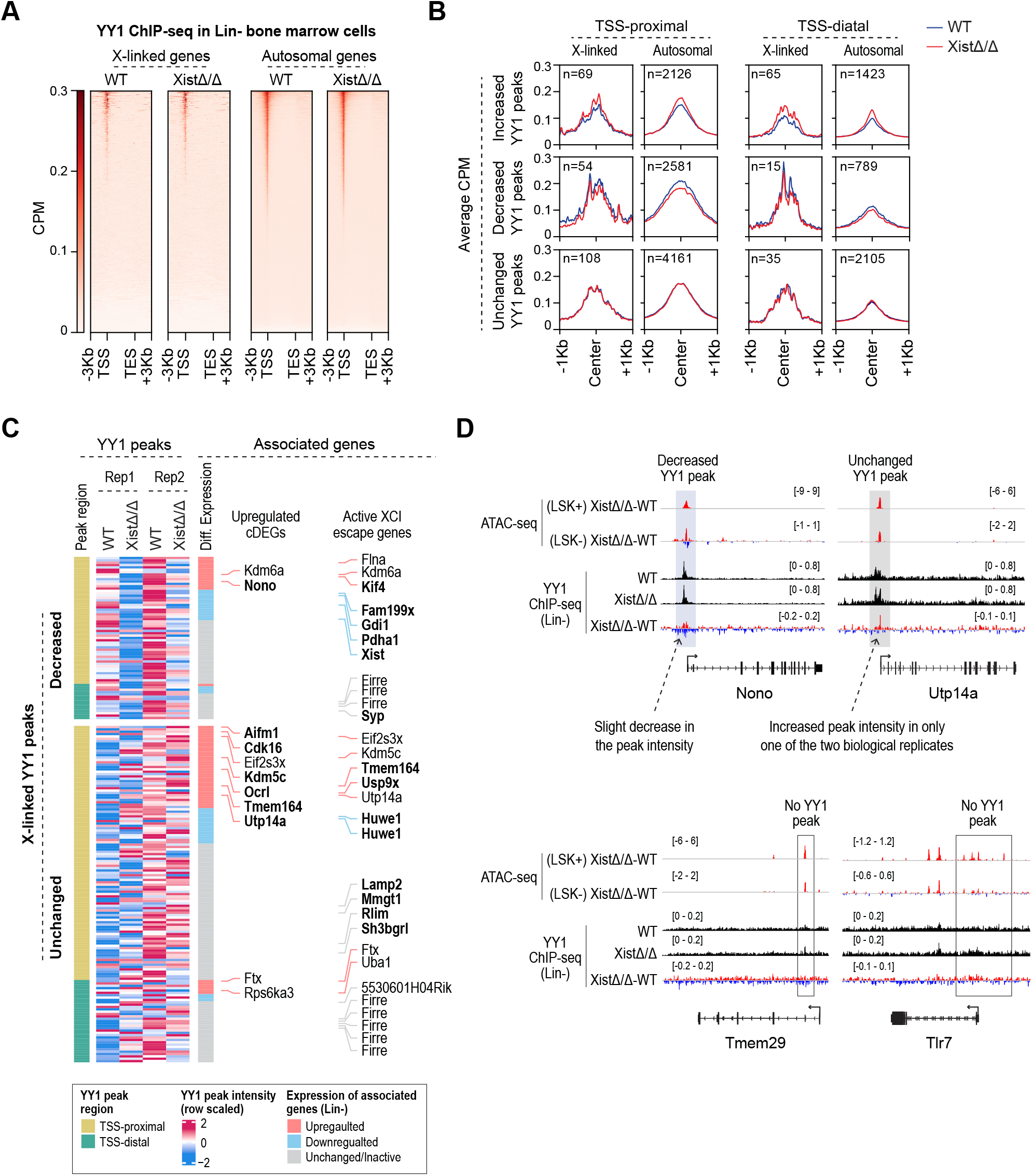
Analysis of YY1 binding and its association with transcription regulation in Xist deficient cells. Related to **Figure 3 and 6.** **(A)** Heatmaps of ChIP-seq identified YY1 binding across +/- 3Kb of all X-linked and autosomal genes in Lin- cells from 3-month-old WT and XistΔ/Δ female mice. CPM, counts per million. **(B)** Metagene profiles show distribution of average YY1 ChIP-seq signal +/- 1Kb from the center of increased, decreased or unchanged YY1 peaks at TSS-proximal and TSS-distal regions on X chromosome and autosomes in WT and XistΔ/Δ Lin- cells. The signal change is larger for increased YY1 peaks than decreased YY1 peaks on the X chromosome. **(C)** Heatmap shows correlation between decreased and unchanged YY1 peaks on the X chromosome, and expression of associated genes at TSS-proximal and TSS-distal regions. Names of X-linked upregulated cDEGs and active mouse XCI escape genes reported by Berletch et al. were indicated. Genes that do not associate with increased YY1 peaks were marked as bold. **(D)** YY1 ChIP-seq and ATAC-seq tracks are shown for four upregulated X-linked DEGs (Tmem29, Tlr7, Nono and Utp14a) that do not associate with increased YY1 binding and ATAC sites. Decreased, unchanged, and no YY1 peak regions are highlighted.

## Notes

### Competing Interest Statement

The authors have declared no competing interest.

